# The DCC receptor regulates astroglial development essential for telencephalic morphogenesis and corpus callosum formation

**DOI:** 10.1101/2020.08.03.233593

**Authors:** Laura Morcom, Ilan Gobius, Ashley P L Marsh, Rodrigo Suárez, Caitlin Bridges, Yunan Ye, Laura R Fenlon, Yvrick Zagar, Amelia M Douglass, Amber-Lee S Donahoo, Thomas Fothergill, Samreen Shaikh, Peter Kozulin, Timothy J Edwards, Helen M Cooper, IRC5 Consortium, Elliott H Sherr, Alain Chédotal, Richard J Leventer, Paul J Lockhart, Linda J Richards

**Affiliations:** The University of Queensland, Queensland Brain Institute, Brisbane, QLD 4072, Australia; Bruce Lefroy Centre for Genetic Health Research, Murdoch Children’s Research Institute, Royal Children’s Hospital, Parkville, VIC 3052, Australia; Department of Paediatrics, University of Melbourne, Parkville, VIC 3052, Australia; Sorbonne Université, INSERM, CNRS, Institut de la Vision, 17 Rue Moreau, 75012 Paris, France; The University of Queensland, Faculty of Medicine, Brisbane, QLD 4072, Australia; Members and Affiliates of the International Research Consortium for the Corpus Callosum and Cerebral Connectivity (IRC5); Departments of Neurology and Pediatrics, Institute of Human Genetics and Weill Institute of Neurosciences, University of California, San Francisco, San Francisco, CA 94143, USA; Neuroscience Research Group, Murdoch Children’s Research Institute, Parkville, VIC 3052, Australia; Department of Neurology, University of Melbourne, Royal Children’s Hospital, Parkville, VIC 3052, Australia; The University of Queensland, School of Biomedical Sciences, Brisbane, QLD 4072, Australia

**Keywords:** Midline zipper glia, astrocyte morphology, agenesis of the corpus callosum, callosal axons, Deleted in colorectal cancer, NTN1, DCC mutations, telencephalic development, interhemispheric fissure remodelling

## Abstract

The forebrain hemispheres are predominantly separated during embryogenesis by the interhemispheric fissure (IHF). Radial astroglia remodel the IHF to form a continuous substrate between the hemispheres for midline crossing of the corpus callosum (CC) and hippocampal commissure (HC). DCC and NTN1 are molecules that have an evolutionarily conserved function in commissural axon guidance. The CC and HC are absent in *Dcc* and *Ntn1* knockout mice, while other commissures are only partially affected, suggesting an additional aetiology in forebrain commissure formation. Here, we find that these molecules play a critical role in regulating astroglial development and IHF remodelling during CC and HC formation. Human subjects with *DCC* mutations display disrupted IHF remodelling associated with CC and HC malformations. Thus, axon guidance molecules such as DCC and NTN1 first regulate the formation of a midline substrate for dorsal commissures prior to their role in regulating axonal growth and guidance across it.

## Introduction

The corpus callosum (CC) is the largest fibre tract in the human brain and is comprised of approximately 200 million axons (Paul et al., 2007; Tomasch, 1954) connecting similar regions between the left and right cerebral hemispheres (Fenlon and Richards, 2015; Fenlon et al., 2017; Suárez et al., 2018). All eutherian mammals have a corpus callosum (Suárez et al., 2014; Suárez, 2017), with malformations or complete absence (agenesis) of the CC occuring in at least 1 in 4000 human live births (Glass et al., 2008). Collectively, these genetically heterogeneous disorders are known as CC dysgenesis, and can result in a wide spectrum of neurological, developmental and cognitive deficits (Brown and Paul, 2019; Edwards et al., 2014; Paul et al., 2007).

During brain development, the callosal tract forms between the two telencephalic hemispheres through a midline region initially separated by the interhemispheric fissure (IHF; Gobius et al., 2016; Rakic and Yakovlev, 1968; Silver et al., 1982). Recently, we demonstrated that remodelling of the IHF tissue by specialised astroglial cells, known as midline zipper glia (MZG), is mediated by FGF8 signalling and subsequent regulation of astrogliogenesis by NFI transcription factors, and is essential to provide a permissive substrate for callosal axons to cross the telencephalic midline (Gobius et al., 2016). MZG are derived from radial glia in the telencephalic hinge, located rostral to the third ventricle. From this ventricular zone, they migrate rostro-dorsally as bipolar cells to the IHF pial surface and transition into multipolar astrocytes. This latter step facilitates their intercalation across the midline and subsequent elimination of the intervening leptomeningeal tissue that comprises the IHF. The MZG thereby fuse the medial septum in a fashion that resembles a ‘zipper’ mechanism (Gobius et al., 2016), which does not occur in naturally acallosal mammals such as monotremes and marsupials (Gobius et al., 2017). Developmental defects in IHF remodelling invariably result in callosal agenesis in mouse models and, strikingly, all 38 individuals in a human cohort with callosal agenesis also displayed aberrant retention of the IHF and an abnormal separation of the medial septum (Gobius et al., 2016). Thus, the remarkably high prevalence of midline defects in human callosal disorders suggests that there are additional determinant genes for IHF remodelling that have not yet been identified. These could include axon guidance genes, which are frequently mutated in humans (and mice) with CC abnormalities (Edwards et al., 2014).

Netrin 1 (NTN1) is a secreted ligand for the deleted in colorectal carcinoma (DCC) receptor, and these molecules function as axon guidance cues in species ranging from *Drosophila* to mammals (Chan et al., 1996; de la Torre et al., 1997; Fazeli et al., 1997; Hedgecock et al., 1990; Keino-Masu et al., 1996; Kolodziej et al., 1996; Serafini et al., 1996). Indeed, NTN1-DCC signalling attracts pioneering callosal axons towards the midline and attenuates chemorepulsive signaling in neocortical callosal axons *ex vivo* to facilitate crossing the midline (Fothergill et al., 2014). Heterozygous and homozygous *DCC* pathogenic variants also result in human callosal dysgenesis at high frequency (Jamuar et al., 2017; Marsh et al., 2018; Marsh et al., 2017) with an estimated incidence of 1 in 14 in unrelated individuals with callosal dysgenesis (Marsh et al., 2017), and *Ntn1* and *Dcc* mouse mutants do not form a CC (Fazeli et al., 1997; Finger et al., 2002; Fothergill et al., 2014; Serafini et al., 1996). Instead of crossing the midline, callosal axons in *Ntn1* and *Dcc* mutant mice form ipsilateral “Probst” bundles that run parallel to the midline (Fazeli et al., 1997; Finger et al., 2002; Fothergill et al., 2014; Ren et al., 2007; Serafini et al., 1996). Together, these results have led to the conclusion that NTN1 and DCC act primarily as axon guidance genes during callosal formation. However, in *Ntn1* and *Dcc* mutant mice, only the CC and hippocampal commissure (HC) are completely absent, while other axon tracts remain intact or are mildly affected (Fazeli et al., 1997; Serafini et al., 1996; Yung et al., 2015), indicating that additional processes might affect the development of the CC and HC in these mice. Moreover, elimination of the leptomeninges, which normally occurs during IHF remodelling (Gobius et al., 2016), is severely disrupted in *Ntn1* mutant mice (Hakanen and Salminen, 2015), further suggesting that NTN1 and its receptor, DCC, may play a hitherto unidentified role in IHF tissue remodelling.

Here, we identify a distinct and developmentally earlier role for NTN1 and DCC signalling during CC formation, involving the regulation of MZG development and subsequent IHF remodelling. We find that IHF remodelling is impaired in both *Ntn1* and *Dcc* mouse mutants, as well as in humans with *DCC* pathogenic variants that also display agenesis of the CC and HC. Moreover, in contrast to the wildtype receptor, these human pathogenic variants of *DCC* are unable to regulate cell morphology. Furthermore, we find that defects in astroglial morphology and migration to the IHF in *Ntn1* and *Dcc* mutant mice prevent MZG intercalation and, therefore, IHF remodelling and midline crossing of commissural axons. Taken together, our findings indicate that pathogenic variants in *NTN1* and *DCC* are most likely to affect human CC and HC development through misregulation of astroglial shape, motility and function during IHF remodelling.

## Results

### *Dcc* signalling is required for IHF remodelling and subsequent CC and HC formation

To re-investigate how *Dcc* and *Ntn1* regulate callosal formation, we first analysed the relationship between the IHF and callosal axon growth during midline development in horizontal sections of *Ntn1* and *Dcc* mutant mice. These mouse mutants include *Dcc* knockout, *Dcc^kanga^* mice, which express a truncated DCC receptor that lacks the P3 intracellular signalling domain, and *Ntn1-lacZ* mutant mice, which express reduced levels of NTN1 protein that subsequently becomes sequestered in intracellular organelles (Fazeli et al., 1997; Finger et al., 2002; Fothergill et al., 2014; Serafini et al., 1996). Immunohistochemistry was performed following commissure formation at embryonic day (E)17 against the axonal marker Gap43 together with pan-Laminin, which labels both leptomeningeal fibroblasts and the basement membrane surrounding the IHF (Figure 1A). This revealed that commissural axons in *Dcc* knockout, *Dcc^kanga^*, and *Ntn1-lacZ* mice remain within the ipsilateral hemisphere and do not form a CC or HC, consistent with previous reports (Fazeli et al., 1997; Finger et al., 2002; Fothergill et al., 2014; Ren et al., 2007; Serafini et al., 1996). We further identified that IHF remodelling had not occurred in *Dcc* knockout, *Dcc^kanga^*, and *Ntn1-lacZ* mice, evidenced by complete retention of the IHF, which separated the majority of the telencephalic midline (Figure 1A). This likely prevented formation of the HC in addition to the CC (Figure 1A). The extent of IHF retention, measured as the ratio of IHF length to total midline length, is significantly larger in *Dcc* and *Ntn1* mutants compared to their wildtype littermates (Table S1; Figure 1A and 1B), but did not differ between mutants (Table S1; Figure 1A and 1B). This suggests that NTN1 and DCC may interact or act in a similar manner to regulate IHF remodelling prior to commissural axon crossing, and that the P3 intracellular domain of DCC is crucial for this function. The brain phenotype of adult *Dcc* knockout and *Ntn1-lacZ* mice was unable to be investigated as these animals die shortly after birth (Fazeli et al., 1997; Finger et al., 2002; Serafini et al., 1996). However, immunohistochemistry for the axonal marker Neurofilament in adult *Dcc^kanga^* mice revealed that the retention of the IHF and absence of the CC and HC persists into adulthood (Table S1; Figures 1B and S1), resembling human congenital callosal agenesis (Edwards et al., 2014; Gobius et al., 2016).

**Figure 1:**
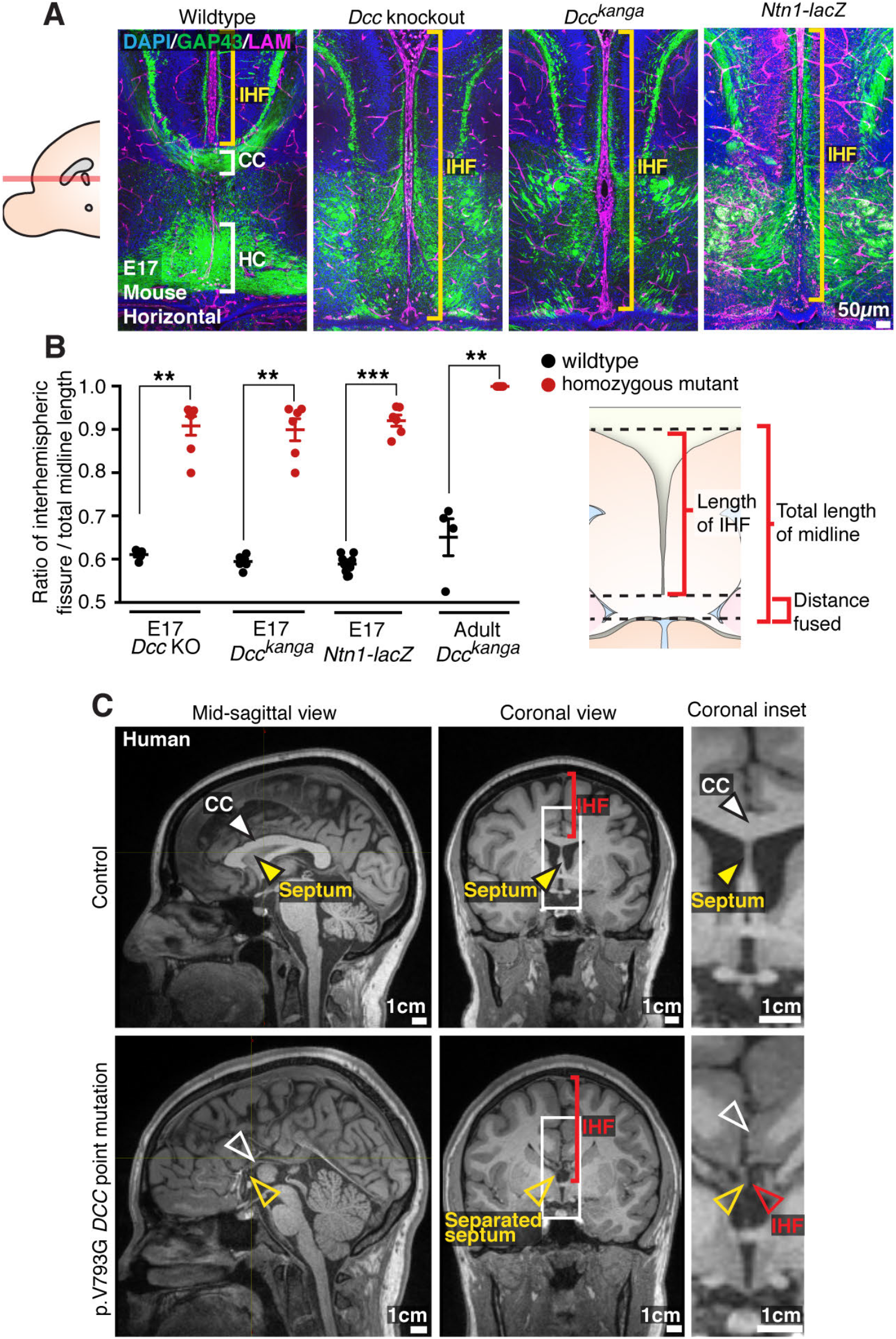
NTN1 and DCC are crucial for remodelling of the IHF, CC and HC formation. (A) Staining for Gap43-positive axons (green) and pan-Laminin (LAM)-positive leptomeninges, and basement membrane (magenta) in wildtype, *Dcc* knockout and *Dcc^kanga^* mice at E17, indicate midline formation or absence of the CC and HC (white brackets) and extent of the IHF (yellow brackets). (B) The ratio of IHF length over the total midline length with schema. (C) T1-weighted MR images of a control subject compared with an individual with a DCC mutation demonstrate the presence or absence of the CC (white arrowheads) and extent of the IHF (red arrowheads and brackets) within the septum (yellow arrowheads). Graph represents mean ± SEM. Statistics by Mann-Whitney test: **p < 0.01, ***p < 0.001. See related Figure S1 and Table S1.

We previously reported that humans carrying *DCC* pathogenic variants develop dysgenesis of the CC with incomplete penetrance (Marsh et al., 2017). T1-weighted MRI of four individuals from two unrelated families carrying missense pathogenic variants in *DCC* (p.Val793Gly affecting fibronectin type III-like domain 4 of *DCC* and p.Met1217Val; p.Ala1250Thr affecting the cytoplasmic domain of *DCC*; Figure 8A; Marsh et al., 2017), revealed in all cases that the complete absence of the CC was associated with aberrant retention of the IHF and an unfused septum (Figure 1C). Importantly, these individuals were also previously reported to lack a HC (Marsh et al., 2017), suggesting a defect in IHF remodelling may also impact HC development. Since IHF remodelling is required for subsequent callosal axon crossing (Gobius et al., 2016), these results collectively suggest that the underlying cause of callosal agenesis in *Ntn1* and *Dcc* mutant mice and in humans with *DCC* mutations is a failure of IHF remodelling.

### DCC and NTN1 are expressed by MZG cells throughout interhemispheric remodelling

We previously demonstrated that DCC is expressed on axons of the CC, HC and the fornix during midline development, while NTN1 is expressed at the telencephalic midline, within the indusium griseum and the septum but not within callosal axons themselves (Fothergill et al., 2014; Shu et al., 2000). Since our analysis of *Ntn1* and *Dcc* mutant mice revealed that these genes are necessary for IHF remodelling, we then investigated whether they are expressed by the MZG, which mediate IHF remodelling (Gobius et al., 2016). MZG arise in the telencephalic hinge, a region in the septal midline caudal to the IHF and rostral to the third ventricle. Radial glia within the telencephalic hinge are attached to both the third ventricle and the IHF and mature into MZG as they undergo somal translocation to the IHF between E12 and E16 in mice (Gobius et al., 2016). Fluorescent *in situ* hybridization for *Dcc* and *Ntn1* transcripts, combined with immunohistochemistry for the MZG marker Glast (Gobius et al., 2016), revealed *Dcc* and *Ntn1* expression in radial MZG progenitor cells within the telencephalic hinge at E12 and E15 (Figure 2B-2D, 2F and 2H; Figure S2H-J), and in MZG migrating to the IHF at E15 (Figure 2F and 2H). Furthermore, *Dcc* was expressed in Glast-positive radial glia within the septum but not in the neocortex (Figure S2A-C). DCC protein can be identified on Glast-positive processes of radial glia attached to the IHF (Figure 2G), which are adjacent to Gap43-positive axons traversing the midline region that also express DCC (Figure S2E). Following IHF remodelling at E17, mature Gfap-positive/Sox9-positive multipolar MZG cells (Gobius et al., 2016; Sun et al., 2017) and Glast-positive MZG cells within the telencephalic hinge continue to express DCC (Figure 2J-2L). A comparison of DCC immunohistochemistry in wildtype and *Dcc* knockout mice confirmed that the antibody specifically recognised DCC protein within both commissural axons and MZG cells (Figure S2K). Importantly, we did not observe specific staining for either *Dcc or Ntn1* mRNA within the IHF (including the leptomeninges) at any stage analysed (Figures 2 and S2).

**Figure 2:**
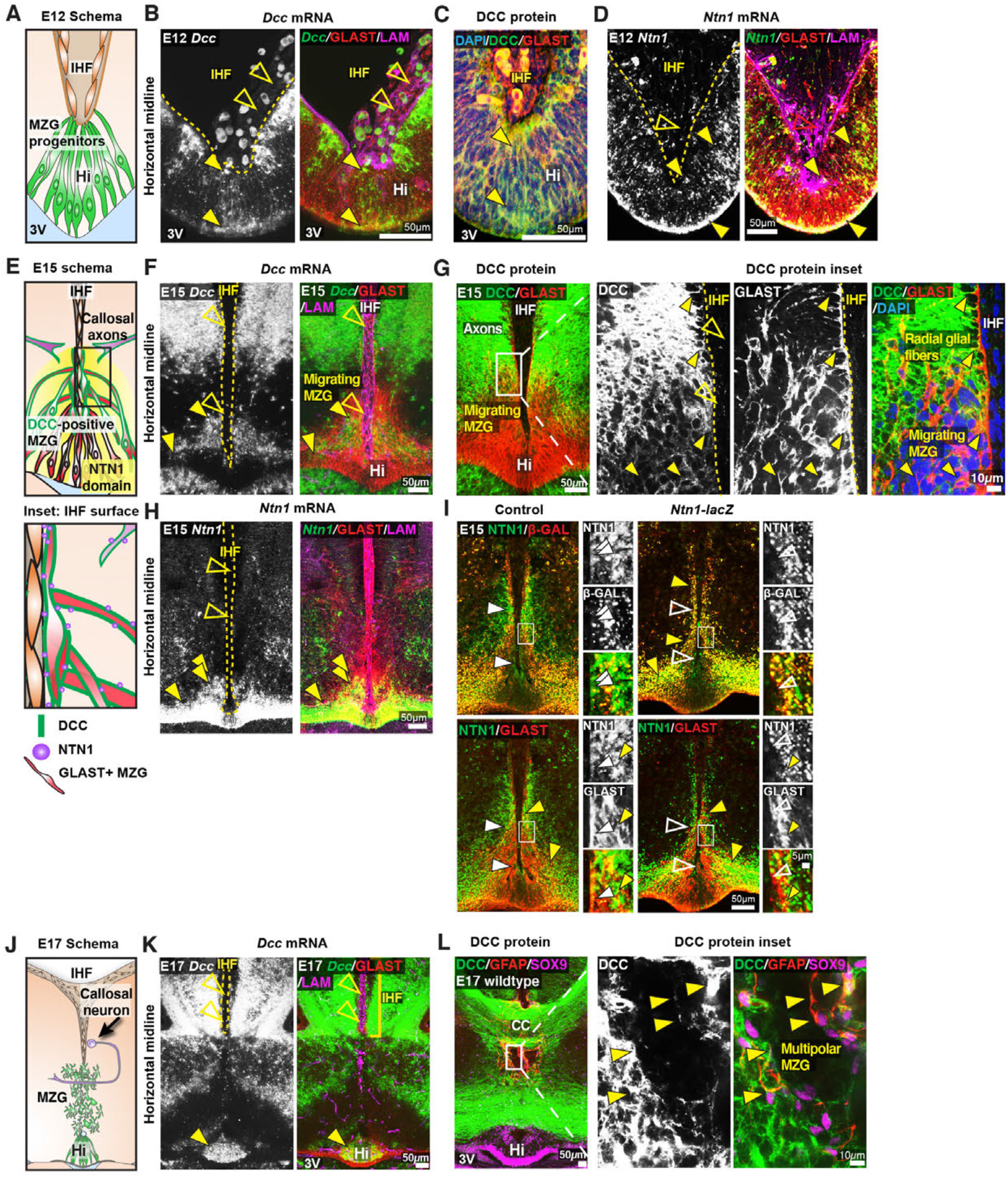
DCC and NTN1 are expressed in MZG and MZG progenitors. (A, E, and J) Schematics depicting the cellular composition of the ventral telencephalic midline at E12, E15 and E17. (B, F and K) *Dcc* mRNA (green), Glast-positive glia (red), and pan-laminin (LAM)-positive leptomeninges and basement membrane (magenta) in E12, E15, and E17 wildtype mice reveal *Dcc*-positive/Glast-positive glial fibers (yellow arrowheads) and absence of *Dcc* within the IHF (open yellow arrowheads). (C and G) DCC protein (green) and Glast protein (red) at E12 and E15 in wildtype mice reveal DCC-positive/Glast-positive glial fibers (yellow arrowheads) and absence of DCC within the IHF (open yellow arrowheads). (D and H) *Ntn1* mRNA (green), Glast (red) and pan-LAM (magenta) in E12 and E15 wildtype mice show *Ntn1*-positive/Glast-positive glial fibers (yellow arrowheads) and absence of *Ntn1* within the IHF (open yellow arrowheads). (E inset) Schema of DCC and NTN1 expression at the E15 IHF surface, based on the results from F-I and Figure S2. (I) NTN1 (green) and Glast (red) or β-galactosidase (β-GAL; red) immunolabelling in E15 control and *Ntn1-lacZ* mice identify regions of NTN1 staining present in control heterozygotes and absent in homozygous *Ntn1-lacZ* mice (white arrowheads) and NTN1-/β-GAL-positive puncta located in Glast-positive glia (yellow arrowheads), with insets. (L) DCC protein (green), glial-specific nuclear marker SOX9 (magenta) and mature astroglial marker (GFAP) in E17 wildtype mice identify DCC-positive/GFAP-positive/SOX9-positive glia (yellow arrowheads). 3V = third ventricle, Hi = telencephalic hinge, See related Figure S2.

Since NTN1 is a secreted cue (Kennedy et al., 1994; Sun et al., 2011), we investigated which cells express NTN1, and where secreted NTN1 may be deposited, by comparing patterns of immunohistochemistry for β-galactosidase (β-gal) and NTN1 antibodies in heterozygous and homozygous *Ntn1-lacZ* mutants, in which NTN1 is fused to a β-gal and trapped in intracellular compartments (Serafini et al., 1996). NTN1/βgal-positive puncta were enriched in Glast-positive MZG cells in *Ntn1-lacZ* mice (Figure 2I). Furthermore, we identified NTN1 protein on the IHF basement membrane (Figures 2I and S2G), on growing commissural axons (Figure S2G), and on MZG membranes in control heterozygotes, but not in *Ntn1-lacZ* homozygous mutant mice (Figure 2I). Therefore, MZG cells produce and secrete NTN1 that becomes deposited on the basement membrane of the IHF, on commissural axons, and on MZG cell processes in the region of initial IHF remodelling (Figure 2E). Collectively, our results demonstrate that both *Ntn1* and *Dcc* are expressed by MZG, and suggest that autocrine NTN1-DCC signalling may regulate MZG development and subsequent IHF remodelling.

### *Dcc* signalling regulates MZG cell morphology and process organisation prior to IHF remodelling

Two key steps in IHF remodelling are the somal translocation of radial MZG progenitors to the IHF, and their subsequent transition into multipolar MZG cells that intercalate across the midline (Gobius et al., 2016). As both NTN1 and DCC are expressed by MZG, we next asked whether these molecules regulate MZG development. Immunohistochemistry for Nestin and Glast, which are markers of radial MZG, revealed distinct differences in MZG development in *Dcc^kanga^* mice from E14 onward, but not in radial MZG progenitors at E13 (Figure S3A). In wildtype mice, the endfeet and cell bodies of radial Glast-positive MZG cells are evenly distributed along the medial septum and adjacent to the pial surface of the IHF (Figures 3B, 3D, 4A and 4C). However, in *Dcc^kanga^* mutants, radial MZG accumulate at the base of the IHF (Figure 3A-D). Furthermore, long radial Nestin-positive MZG processes extending from the ventricular zone to the rostral-most pial surface of the IHF are noticeably absent from *Dcc^kanga^* mutants, and instead, Nestin-positive *Dcc^kanga^* processes cluster close to the rostral IHF pial surface and appear disorganised (Figures 3A, 3C, 3C’, 3E and S3B-D). These abnormalities were further quantified as a significant increase in fluorescence intensity of Glast staining within the base of the IHF, and a concomitant decrease in the region 150-200 µm distant from the IHF base in *Dcc^kanga^* mutants, compared to their wildtype littermates at E14 (Table S1; Figure 3B, 3B’ and 3G). Just prior to IHF remodelling at E15, there was an overall decrease in Glast-positive radial MZG processes in *Dcc^kanga^* mutants (Figure 3C, 3H and Table S1). While there was no difference in fluorescence intensity of Glast-positive radial MZG processes one day later at E16, *Dcc^kanga^* MZG processes continued to display irregular morphology and failed to intercalate across the IHF (Figure 3E, 3F, 3I and Table S1). Interestingly, we identified a similar defect in the distribution of Glast-positive MZG processes in *Ntn1-lacZ* mutant mice at E15 (Figure 3K). These results suggest that both DCC and NTN1 are required for the correct morphology and distribution of MZG processes prior to IHF remodelling.

**Figure 3:**
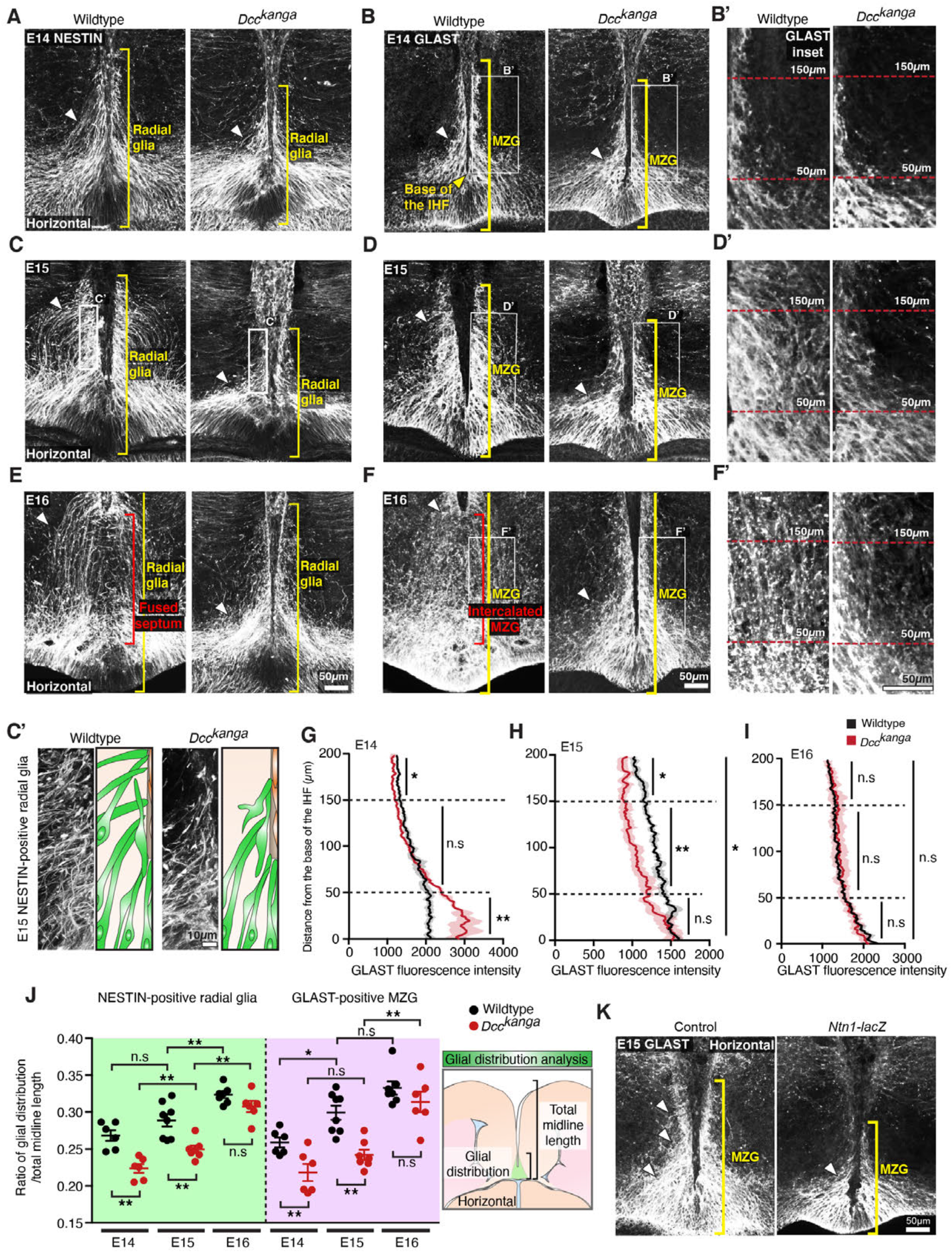
NTN1 and DCC regulate MZG morphology and spatial distribution. Nestin-positive radial glia (white; A, C and E) and Glast-positive glia (white; B, D, F and K) in E14 - E16 *Dcc^kanga^* mice (A-F) and E15 *Ntn1-LacZ* mice (K) demonstrate the distribution of glial processes along the IHF surface (yellow brackets) and lateral to the IHF (white arrowheads) with insets (C’, B’, D’ and F’). The mean fluorescence intensity of Glast staining between wildtype and *Dcc^kanga^* mice at E14 (G), E15 (H) and E16 (I) based on the results from B, D and F, respectively. (J) The ratio of glial distribution over total midline length, with schema, based on the results from A-F. All graphs represent mean ± SEM. Statistics by Mann-Whitney test (C), or a Two-way ANOVA test with post Sidak’s multiple comparison test (D): n.s = not significant, *p < 0.05, **p < 0.01. See related Figure S3 and Table S1.

**Figure 4:**
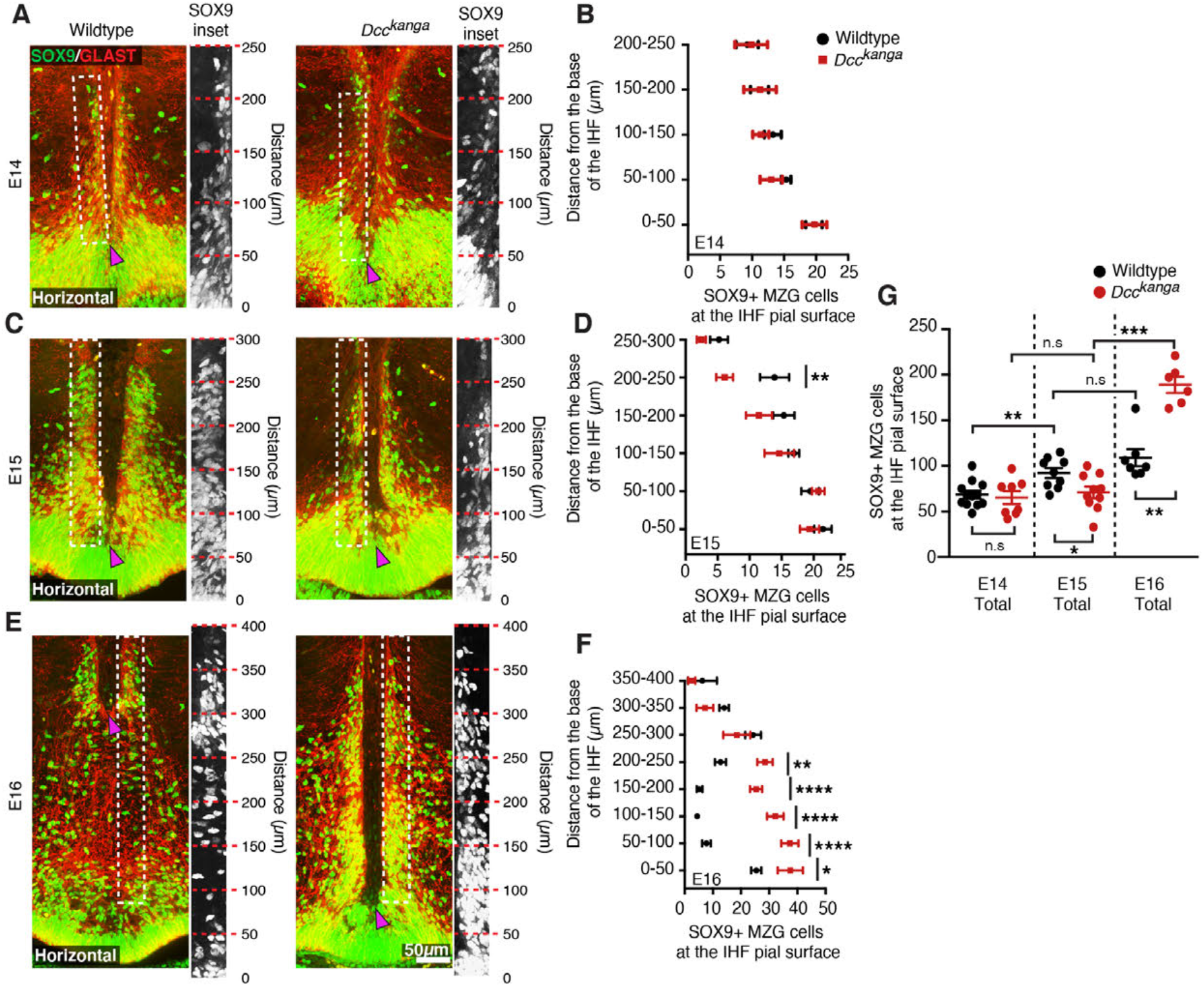
DCC regulates MZG migration to the IHF surface. (A, C, E) Nuclear glial marker SOX9 (green) and MZG marker Glast (red), in E14-E16 *Dcc^kanga^* mice reveal SOX9-positive/Glast-positive MZG at the pial IHF surface (boxed region and insets) above the base of the IHF (magenta arrowhead). (B, D, F, G) Quantification of SOX9-positive/Glast-positive MZG at the IHF pial surface based on the results from A, C and E. All graphs represent mean ± SEM. Statistics by Mann-Whitney test (E) or Two-way ANOVA with post Sidak’s multiple comparison test (B-D): *p < 0.05, **p < 0.01, ***p < 0.001, ****p < 0.0001, n.s = not significant. See related Figure S4 and Table S1.

To further characterise the defect in MZG cell distribution in *Dcc^kanga^* mice, we then measured the maximum rostro-caudal extent to which MZG occupy the IHF pial surface, and normalised this value to the total midline length from E14-E16 (Figure 3A-F and 3J). The distribution of Nestin-positive and Glast-positive MZG along the IHF was significantly decreased at both E14 and E15 in *Dcc^kanga^* mice compared to their wildtype littermates (Figure 3A-D, 3J and Table S1). The attachment of MZG processes to the IHF pial surface is therefore specifically reduced in the rostral region of the IHF prior to IHF remodelling in *Dcc^kanga^* mice. This may impact the directed somal translocation of *Dcc^kanga^* MZG cell bodies and their subsequent distribution along the IHF surface prior to IHF remodelling.

Next, we further investigated whether the aberrant organisation of radial glial processes along the IHF in *Dcc^kanga^* mice was due to a loss of endfoot adhesion to the IHF pial surface. There was no difference in fluorescence intensity of Nestin-positive MZG processes within 5 µm adjacent to the IHF surface between *Dcc^kanga^* and wildtype mice, suggesting comparable attachment of radial glial endfeet to the IHF in both strains (Table S1, Figure S3E and S3G). This was further evidenced by the normal localisation of α- and β-dystroglycan at the pial IHF surface in *Dcc^kanga^* mice, where these molecules form crucial adhesions between radial glial endfeet and the extracellular matrix (Myshrall et al., 2012; Table S1; Figure S3-D). Moreover, molecules that normally maintain the bipolar morphology of radial glia, such as β-catenin and N-cadherin, were also expressed normally within *Dcc^kanga^* Nestin-positive radial glia, but adenomatous polyposis coli (APC) was instead significantly reduced in *Dcc^kanga^* Nestin-positive radial glial endfeet (Table S1, Figure S3D and S3E; Yokota et al., 2009). APC regulates the growth and extension of basal radial glial processes and cell polarity of radial glia and migrating astrocytes (Etienne-Manneville and Hall, 2003; Yokota et al., 2009). Thus, reduced localisation of APC within *Dcc^kanga^* radial glial basal endfeet may indicate perturbed regulation of cell process growth and/or cell polarity. Therefore, *Dcc^kanga^* Nestin-positive radial glia display reduced elongation and reduced occupation of the pial IHF surface compared to wildtype radial progenitors of MZG. Collectively, these results suggest that DCC is not required for radial MZG to adhere to the IHF, but instead regulates the morphology and organisation of radial MZG processes along the pial IHF surface prior to IHF remodelling.

### *Dcc* signalling regulates MZG somal translocation to the IHF prior to IHF remodelling

To determine if the aberrant morphology and organisation of radial glial processes observed in *Dcc^kanga^* mice affects the subsequent distribution of translocated MZG cell bodies at the IHF surface, immunohistochemistry for glial markers Sox9 and Glast was performed in E14-E16 *Dcc^kanga^* mice. Wildtype MZG undergo substantial somal translocation to the IHF between E14 and E15 (Gobius et al., 2016; Table S1, Figure 4A, 4C and 4G). In contrast, *Dcc^kanga^* mice showed reduced somal translocation to the IHF (Table S1; Figure 4A, 4C and 4G), with significantly fewer MZG cells at the IHF pial surface by E15 in *Dcc^kanga^* compared to wildtype mice (Table S1; Figure 4B and 4G). When binned along the rostro-caudal axis, we found a significant reduction in the number of cell bodies reaching the rostral IHF pial surface in E15 *Dcc^kanga^* mice (200-250 µm; Table S1 and Figure 4C-D). Since MZG progenitors somal translocate toward their basal process attached to the IHF (Gobius et al., 2016), our results suggest that the lack of radial MZG processes occupying the rostral E14 IHF surface in *Dcc^kanga^* mice results in a decrease of MZG cell bodies present in the corresponding region one day later. There was, however, a significant increase in MZG cell bodies present at the IHF pial surface between E15 and E16 in *Dcc^kanga^* mice (Table S1, Figure 4C, 4E and 4G). This suggests that MZG migration towards the IHF is delayed but does eventually occur in *Dcc^kanga^* mice, albeit after IHF remodelling would normally have been initiated. Furthermore, despite DCC having been previously implicated in regulating cell proliferation and cell death (Arakawa, 2004; Llambi et al., 2001; Mehlen et al., 1998), cell birth-dating, differentiation and apoptosis experiments did not reveal any significant differences between the MZG of *Dcc^kanga^* and wildtype mice (Table S1 and Figure S4). Taken together, these results suggest that the irregular morphology and distribution of radial *Dcc^kanga^* MZG processes is associated with delayed somal translocation of MZG to the IHF surface, and may prevent the initiation of IHF remodelling.

Radial glia in the corticoseptal boundary detach from the pial surface and cluster their processes to form a triangular group of cells known as the glial wedge, while other radial glia in this region translocate their soma to the IHF (similar to MZG cells), where they subsequently form the indusium griseum glia (Shu and Richards, 2001; Smith et al., 2006). We investigated whether DCC also regulates the development of these glial populations, which secrete axon guidance molecules during corpus callosum formation (reviewed in Donahoo and Richards, 2009; Gobius and Richards, 2011; Morcom et al., 2016). In *Dcc^kanga^* and *Dcc* knockout mice, the glial wedge was malformed and there was a major reduction in somal translocation of Sox9-positive indusium griseum glia to the IHF surface, which subsequently prevented formation of this glial guidepost cell population (Table S1 and Figure S4G-I). Thus, DCC may play a more general role in regulating the morphological maturation and migration of multiple radial astroglial populations in the developing midline, which are critical for CC formation.

### DCC signalling regulates MZG cell morphology and spatial distribution during IHF remodelling

We previously demonstrated that MZG differentiation is controlled by molecular signaling initiated by the morphogen FGF8 via the Mitogen activated protein kinase (MAPK) pathway to NFI transcription factors A and B (Gobius et al., 2016). Members of this signaling pathway (*Fgf8*, NFIA, NFIB, and p-ERK1/2) were expressed normally in *Dcc^kanga^* MZG compared to MZG in their wildtype littermates at E15 (Figure S5B, S5D-F and Table S1). Further, *Dcc^kanga^* MZG continue to express *Mmp2* mRNA (Figure S5C, D and Table S1), which we previously demonstrated to be expressed during MZG-mediated degradation of the IHF during remodelling (Gobius et al., 2016). Next, we investigated the distribution and maturation of MZG in *Ntn1* and *Dcc* mutant mice at E16 and 17, when wildtype MZG normally differentiate into multipolar astrocytes during IHF remodelling (Gobius et al., 2016). Immunohistochemistry for Nestin, Glast (Figures 3F, 3J) and Gfap (Figure 5A) demonstrated that *Dcc^kanga^*, *Dcc* knockout and *Ntn1-lacZ* MZG remain attached to the caudal IHF pial surface and have not intercalated at stages equivalent to when wildtype MZG have infiltrated and remodelled the IHF (Figures 3F, 3J, 5A and Table S1). DCC-deficient MZG expressed GFAP at comparable levels to wildtype MZG at E17, and demonstrated no precocious expression of GFAP at E15, similar to wildtype MZG (Figure 5A-B, Figure S5A and Table S1). Therefore, DCC-deficient MZG do not mature precociously prior to migration and IHF remodelling, or fail to differentiate during callosal development. However, both *Dcc^kanga^* and *Dcc* knockout mice demonstrated a significant reduction of GFAP-positive glia at E17 in the region where the corpus callosum normally forms in wildtype mice (i.e., 450-700 µm from the third ventricle; Figure 5A-B and Table S1). *Ntn1-lacZ* mice showed a similar trend that did not reach significance (450-700 µm from the third ventricle; Figure 5A,C and Table S1). In line with this, an increased number of MZG cells remained in the caudal midline in both *Dcc* knockout and *Ntn1-lacZ* mice (0-450 µm from the third ventricle in *Dcc* knockout, 0-450 µm in *Ntn1-lacZ*; Figure 5A-C and Table S1), while MZG in *Dcc^kanga^* mice were more variably distributed, similar to controls (0-450 µm; Figure 5A, B and Table S1). Since progressive intercalation of MZG is required for IHF remodelling (Gobius et al., 2016), these results indicate that *Ntn1* and *Dcc* affect IHF remodelling by regulating the morphology and spatial organisation of both radial MZG progenitors and mature MZG, and therefore their ability to intercalate across the IHF, but not their proliferation and adhesion to the pial IHF surface.

**Figure 5:**
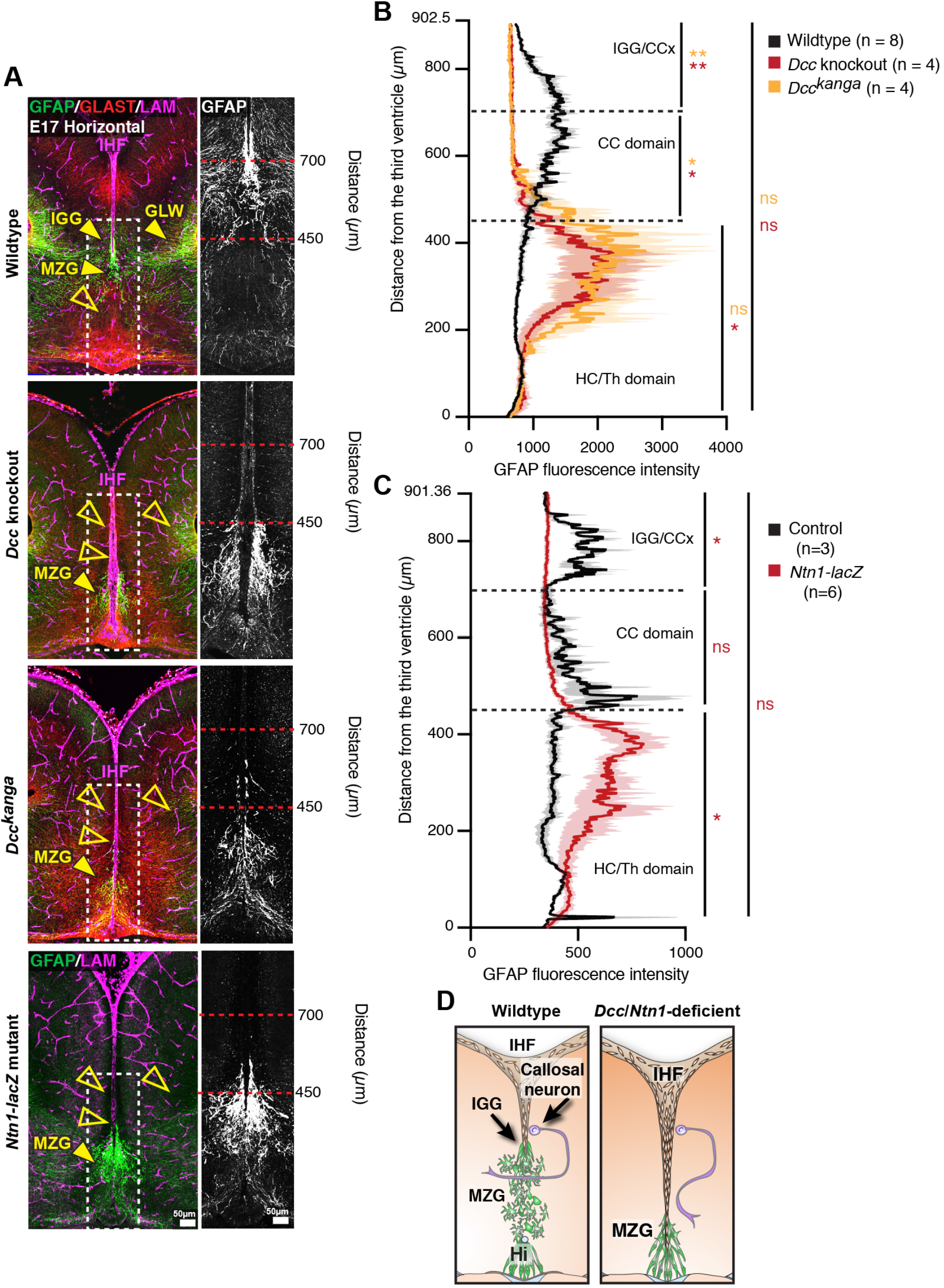
NTN1 and DCC regulate MZG organisation during IHF remodelling. (A) Gfap-positive mature astroglia (green or white in inset), Glast-positive glia (red), and pan-Laminin (LAM)-positive IHF and basement membrane (magenta) in E17 wildtype *Dcc^kanga^*, *Dcc* knockout, and *Ntn1-LacZ* mice. Yellow arrowheads indicate presence (filled) or absence (open) of midline glial populations, the midline zipper glia (MZG), indusium griseum glia (IGG) and glial wedge (GW). Fluorescence intensity of Gfap staining from insets was quantified in B and C. (D) Schema of MZG development, IHF remodelling and CC formation in wildtype mice and mice deficient for NTN1 or DCC. See related Figure S5. All graphs represent mean ± SEM. Statistics by Mann-Whitney test: *p < 0.05, **p < 0.01, n.s = not significant. See related Figure S4 and Table S1.

### Variable DCC knockdown during midline development causes a spectrum of callosal phenotypes

The current and previous results from our laboratory indicate at least two distinct roles for NTN1 and DCC during CC formation: first, they act on astroglia to facilitate remodelling of the IHF, and second, they regulate the pathfinding of callosal axons to the telencephalic midline (Fothergill et al., 2014). To investigate these roles independently, we aimed to disrupt DCC expression specifically within the progenitors of callosal neurons, sparing expression within MZG. We designed two *Dcc*-targeted CRISPR/CAS9 constructs (*Dcc*-CRISPR) and acquired a *Dcc*-targeted shRNA (*Dcc-*shRNA; Zhang et al., 2018) for targeted in utero electroporation into wildtype and *Dcc^kanga^* mice. We reasoned that heterozygous *Dcc^kanga^* mice, which do not display an aberrant phenotype on the C57Bl/6 background (Fothergill et al., 2014), have the potential to express up to 50% non-functional DCC protein that could be further disrupted using *Dcc*-CRISPR or *Dcc-*shRNA techniques. In utero electroporation of either the combined DCC-CRISPR constructs or the *Dcc-*shRNA construct with a pCAG-TdTomato reporter into the E13 cingulate cortex labelled callosal axons that had reached the contralateral hemisphere by E18 (Figure S6A). In all cases, the CC formed normally and the IHF had been remodelled (Figure S6A). However, quantification of DCC expression revealed only a small, significant reduction in DCC protein in heterozygous *Dcc^kanga^* mice electroporated with *Dcc*-shRNA (average DCC expression reduced to 93.06% compared to the non-electroporated hemisphere; Figure S6A-C and Table S1). Thus, we were unable to deplete DCC expression sufficiently in callosal axons to elicit a phenotype. In order to knockout DCC more robustly in the cortex, we crossed *Dcc^flox/flox^* mice (Krimpenfort et al., 2012) with mice carrying *Emx1^iCre^* (Kessaris et al., 2006), and the *tdTomato^flox_stop^* reporter allele (Madisen et al., 2010). These *Dcc* cKO mice were expected to have reduced DCC expression in callosal axons but not in MZG.

At birth, we observed a spectrum of callosal phenotypes in *Dcc* cKO mice, including complete callosal absence (4/12 mice), partial CC absence (5/12 mice), and a normal CC that was comparable to control mice, which do not express *Emx1^iCre^* (3/12 mice) based on rostral-caudal CC length across ventral, middle and dorsal horizontal sections (Figure 6A and 6F). Unexpectedly, we found the IHF was significantly retained across *Dcc* cKO mice, indicating that IHF remodelling had not been completed (Figure 6A, 6D-E, and Table S1). The severity of callosal agenesis was associated with the extent to which the IHF had been remodelled; complete callosal agenesis *Dcc* cKO mice demonstrated the most severe retention of the IHF, encompassing the majority of the telencephalic midline, while partial callosal agenesis and even full CC *Dcc* cKO mice demonstrated a retention of the rostral IHF (Figure 6A, 6E and Table S1). Moreover, in partial callosal agenesis and full CC *Dcc* cKO mice that demonstrated partial retention of the rostral IHF, callosal axons often crossed the midline more caudal in a region where the IHF had been remodelled compared to control mice (see corpus callosum remnant or CCR in Figure 6A). This suggests that in the absence of their normal substrate, callosal axons are able to adapt and cross the midline in a region where the substrate is available. These results were reflected by a significant increase in the rostro-caudal depth of the partial or full CC in *Dcc* cKO mice (Figure 6A, 6E and Table S1). In *Dcc* cKO mice with complete callosal agenesis, callosal axons were unable to cross the midline and accumulated adjacent to the IHF that had not been remodelled (Figure 6A), similar to *Dcc^kanga^* and *Dcc* knockout mice. The HC was significantly reduced in the majority of animals (red arrows in Figure 6A and 6B; Figure 6F and Table S1) and was absent in one *Dcc* cKO mouse, suggesting that IHF remodelling also influences HC formation in mice. These results demonstrate that DCC regulates the extent of IHF remodelling throughout the telencephalic midline. The retention of the IHF in these mice was unexpected; we had instead expected that reduced DCC expression in the cortex would cause callosal axon misguidance with normal formation of an interhemispheric substrate. Instead, our results suggest that DCC primarily regulates the formation of the interhemispheric substrate either indirectly via its expression in axons or directly if *Emx1^iCre^* is expressed in the ventrally located MZG progenitors.

**Figure 6:**
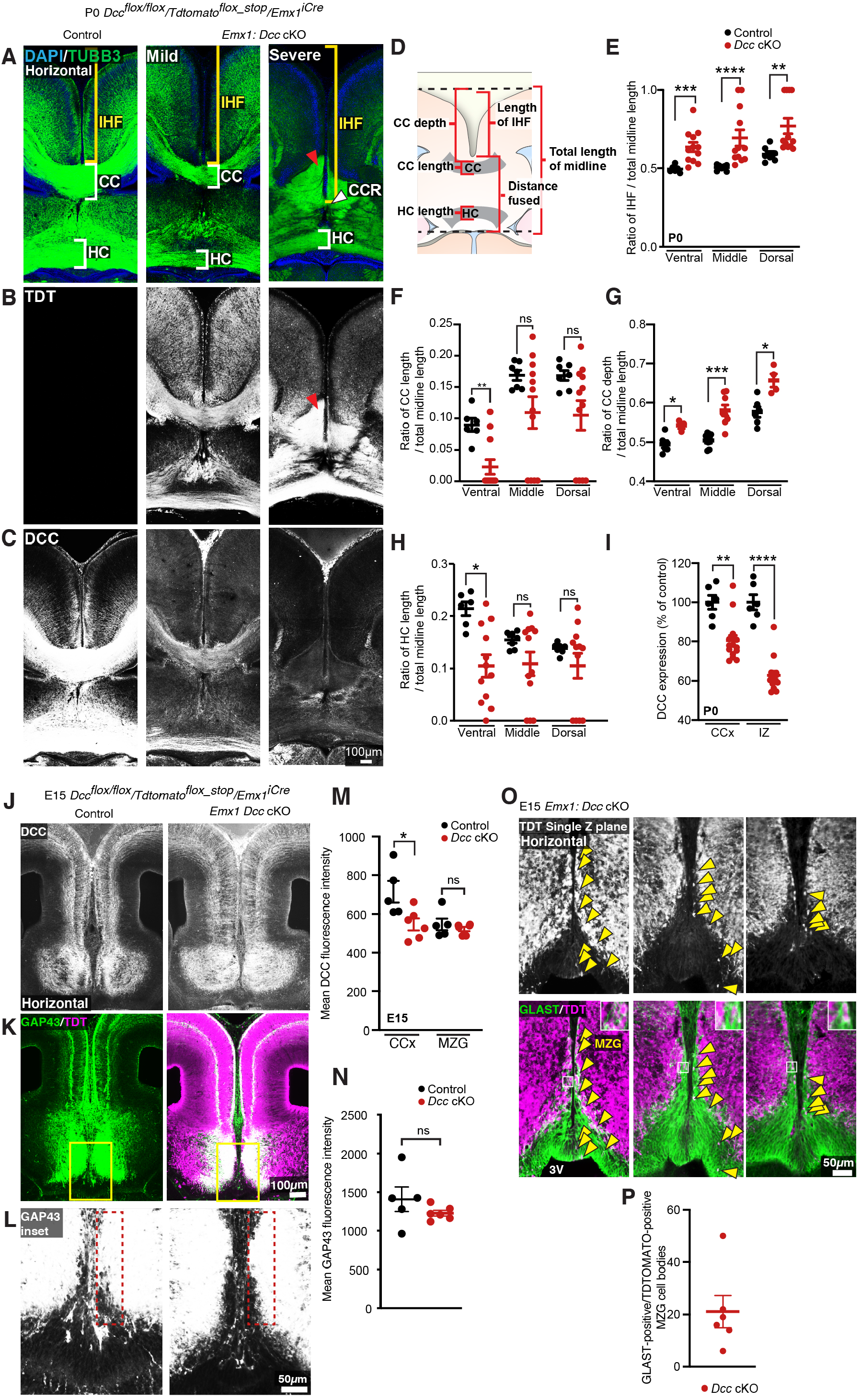
Conditional knockdown of DCC within EMX1 cells causes a spectrum of callosal phenotypes. (A) Axonal marker TUBB3 (green), (B) TDT (white) or (C) DCC (white) in P0 *Dcc* cKO demonstrate a spectrum of callosal and IHF remodelling phenotypes and a reduction in DCC expression within mice expressing *Emx1^iCre^.* The CC or CC remnant (CCR) and HC are indicated with white brackets or white arrowheads, and the IHF is indicated with yellow brackets. Red arrowheads indicate axon bundles that have not crossed the midline. (D) Schema of measurements taken for quantification shown in C-E. (E) Quantification of the ratio of IHF length normalised to total telencephalic midline length measured for P0 *Dcc* cKO mice. (F and G) Quantification of CC length (F) and depth (G) normalised to the total telencephalic midline length in P0 *Dcc* cKO mice. (H) Quantification of HC length normalised to the total telencephalic midline length in P0 *Dcc* cKO mice. (I) Quantification of DCC expression measured from the cingulate cortex (CCx) and intermediate zone (IZ) of *Dcc* cKO mice. (J) DCC (white), (K and L) axonal marker GAP43 (green or white, insets) and TDT (magenta) in E15 *Dcc* cKO mice, with quantification of mean DCC fluorescence in (M), and quantification of mean GAP43 fluorescence within 50 µm from the IHF (dotted red lines) in (N). (O) TDT (white or magenta) and glial marker GLAST (green) in E15 *Dcc* cKO with insets and yellow arrowheads indicating GLAST-positive/TDT-positive MZG, and quantified in P. All graphs represent mean ± SEM. Statistics by Mann-Whitney test or unpaired t test: *p < 0.05, **p < 0.01, ***p < 0.001, ****p < 0.0001, n.s = not significant. See related Figure S5 and Table S1.

We first explored whether loss of DCC expression within callosal axons might cause prior callosal axon misguidance that could indirectly impact IHF remodelling. We found that DCC expression was significantly reduced in the cingulate cortex and adjacent intermediate zone in the majority of P0 *Dcc* cKO mice (mean expression reduced to 80.4% and 62.9% respectively; Figure 6A, 6I and Table S1), and within E15 *Dcc* cKO mice (mean DCC expression in the cingulate cortex reduced to 76.6% in *Dcc* cKO; Figure 6J, 6M and Table S1), as expected. Surprisingly, we found that TDTOMATO-positive/GAP43-positive axons, which had reduced DCC expression, approached the interhemispheric midline in *Dcc* cKO mice, similar to their cre-negative littermates (Figure 6K-L, 6N and Table S1). This suggests that axons with reduced, but not entirely eliminated DCC expression, approach the midline adjacent to the IHF in a timely and spatially appropriate manner, and are unable to cross the midline in regions where the IHF is not remodelled in *Dcc* cKO mice.

Next, we investigated whether MZG cells may be unexpectedly affected in *Dcc* cKO mice. Cre activity, as measured by TDTOMATO expression, was widespread in cells throughout the telencephalic midline, including septal cells and HC axons, resulting in reduced DCC expression in multiple cell types (Figure 6B). Mean DCC expression within the telencephalic hinge was comparable between *Dcc* cKO mice and their cre littermates (Figure 6J, 6M and Table S1), but we also observed TDTOMATO-positive/GLAST-positive MZG cell bodies within the telencephalic hinge, and at the IHF surface in *Dcc* cKO mice (Figure 6O-P and Table S1). This suggests the potential for *Dcc* knockdown in a subset of MZG cells within *Dcc* cKO mice, which may fail to intercalate across the IHF, possibly causing the IHF remodelling defect observed in P0 *Dcc* cKO mice. However, unlike *Dcc^kanga^* mice, we were unable to find a significant population difference in the distribution of GLAST-positive MZG between *Dcc* cKO mice and their cre-negative littermates, at the level of DCC knockdown observed in this model (Figure S6D-E and Table S1). Thus, the variable callosal and IHF remodelling phenotypes observed in *Dcc* cKO mice likely arise from varying degrees of DCC knockdown in these models due to the mosaic expression of *Emx1^iCre^* within MZG. In order to further explore the role of DCC and the impact of human *DCC* mutations on the behavior of astroglia, we next investigated the function of human *DCC* mutations using in vitro assays.

### NTN1-DCC signalling promotes cytoskeletal remodelling of astroglia and neural progenitors

Our results suggest that NTN1 and DCC may have important functions in the morphological development of radial glia more broadly. We established *in vitro* assays to test the function of NTN1-DCC signalling and DCC mutant receptors in regulating the morphology of astroglial-like cells. Such assays can also be used to examine human variants of DCC pathogenic mutations (see next section). To develop these assays, we employed N2A neuroblast cells, which display neural progenitor properties (Augusti-Tocco and Sato., 1969; Shea et al., 1985), as well as U251 glioma cells, which express astroglial markers and display invasive capacity (Zhang et al., 2013) similar to MZG cells. Importantly, endogenous DCC has previously been demonstrated to render several glioma cell lines migratory in response to a gradient of NTN1 as a chemoattractant (Jarjour et al., 2011). Both cell lines were transfected with either full-length DCC fused to a TDTOMATO reporter (pCAG-DCC:TDTOMATO) to express wildtype DCC, or a membrane-targeted TDTOMATO reporter (pCAG-H2B-GFP-2A-Myr-TDTOMATO) as a control and stimulated with NTN1. Moreover, we transfected U251 cells with a DCC^kanga^ construct (pCAG-DCC^kanga^:TDTOMATO), to test whether the P3 domain was critical for NTN1-DCC signalling effects on cell morphology.

Expression of DCC:TDTOMATO in U251 cells in the absence of ligand (vehicle alone) promoted cell spreading and elongation, reflected by a significant increase in average cell area and cell perimeter, and a significant decrease in cell circularity compared to control (Table S1 and Figure 7A-F). This effect was not observed following expression of the DCC^kanga^ construct alone (Table S1 and Figure 7A-F), suggesting that the P3 domain of DCC is critical for inducing changes in glial cell shape. We further confirmed that these morphological changes were due to the presence of the coding region of wildtype DCC, by comparing to cells transfected with plasmids where DCC had been excised and only the TDTOMATO remained (pCAG-TDTOMATO; Table S1 and Figure S7A and S67B). A similar effect was observed following DCC overexpression in N2A cells, which also registered a significant increase in average cell area and cell perimeter, and decrease in cell circularity, compared to controls (Table S1, Figure 7G-I, and Figure S7G and H), further indicating similar effects on cell morphology in glial and neural progenitor lineages. Interestingly, application of NTN1 did not affect cell shape following DCC expression in either cell line (Table S1 and Figure 7A-F), suggesting that endogenous NTN1, which is known to be expressed by U251 cells (Chen et al., 2017), may be sufficient for activation of DCC:TDTOMATO receptors, or that NTN1 is not required for this effect. Typical features of DCC-expressing cells with or without bath application of NTN1 included actin-rich regions resembling filopodia, lamellipodia, and membrane ruffling in U251 cells, while only filopodia were highly abundant in DCC:TDTOMATO-expressing N2A cells; all of these features were rarely observed in control cells from both cell lines (Figure 7A, G). This suggests that DCC signalling promotes remodelling of the actin cytoskeleton in glioma and neuroblast cells in a similar manner to neurons and oligodendrocytes (Rajasekharan et al., 2009; Shekarabi and Kennedy, 2002).

**Figure 7:**
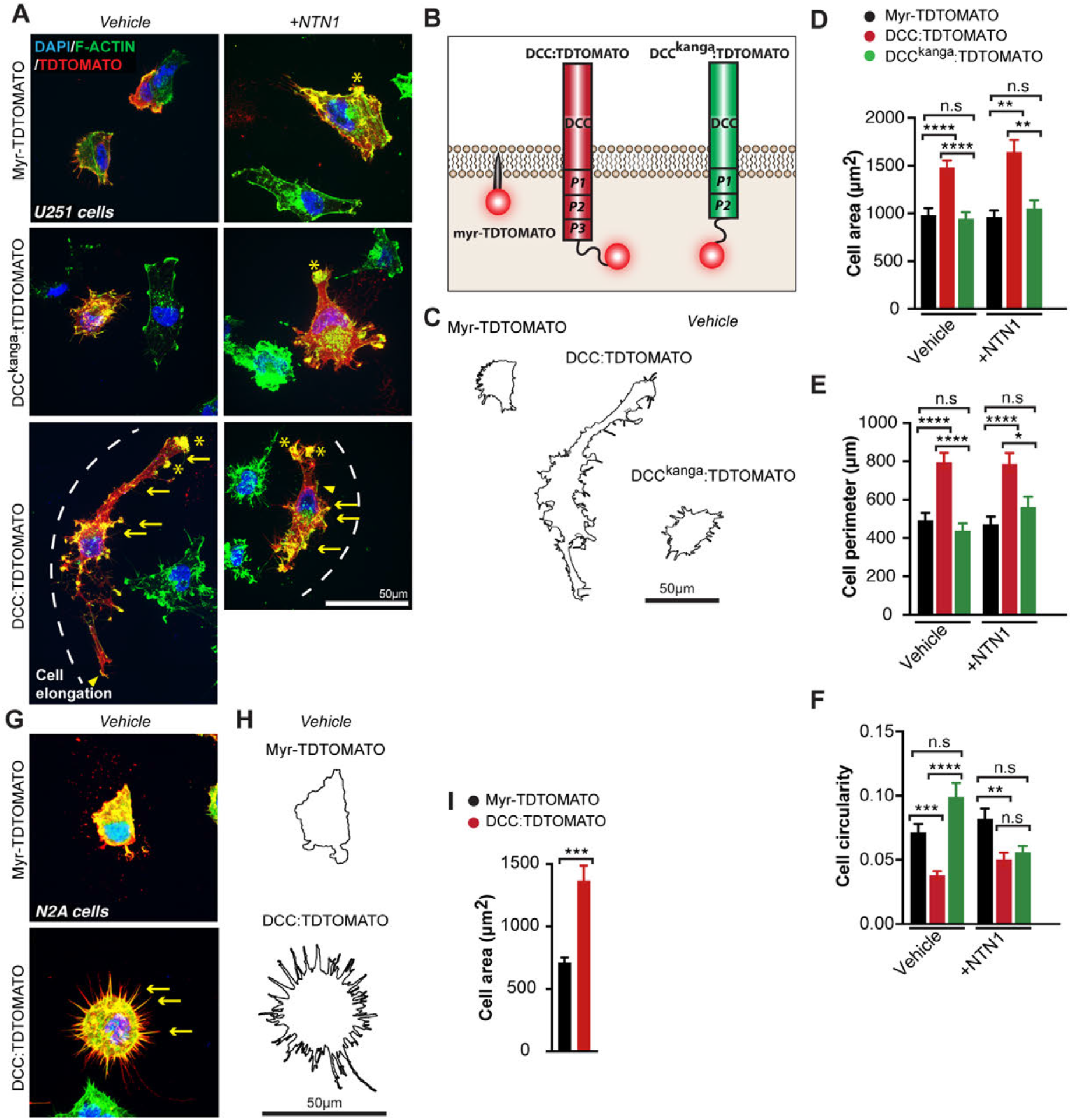
NTN1-DCC signalling promotes cytoskeletal remodelling of astroglia. (A, G) Representative images of U251 glioblastoma cells (A) and N2A cells (G) immunolabelled for TDTOMATO (red), and F-actin (green) following transfection with plasmids encoding Myr-TDTOMATO, DCC:TDTOMATO, or DCC^kanga^:TDTOMATO demonstrating the presence of actin-rich regions resembling filopodia (yellow arrows), lamellipodia (yellow arrowheads) and membrane ruffles (yellow asterisks) with/without stimulation with recombinant mouse NTN1 protein. (B) Schema of predicted structure of proteins on the cell membrane encoded by the plasmids expressed in cells from A and G. (C and H) Outline of cell perimeter generated from images in A and G respectively. (D-F and I) Quantification of the area, perimeter and circularity of cells represented in A and G. Graphs represent mean ± SEM. Statistics by Kruskal-Wallis test for multiple comparisons: n.s = not significant, *p < 0.05, **p < 0.01, ***p < 0.001, ****p<0.0001. See related Figure S6 and Table S1.

### Humans with agenesis of the CC carry loss-of-function pathogenic variants in *DCC* that are unable to modulate cell shape

Having established that DCC signalling rearranges the cytoskeleton of astroglial-like cells, and that the P3 domain of DCC is crucial for this function, we next investigated whether *DCC* mutant receptors from humans with dysgenesis of the CC affected this function. Site directed mutagenesis was performed to introduce missense mutations into the pCAG-DCC:TDTOMATO expression vector in order to model mutated *DCC* receptors found in six families with previously reported cases of complete or partial agenesis of the CC (p.Met743Leu, p.Val754Met, p.Ala893Thr, p.Val793Gly, p.Gly805Glu, p.Met1217Val;p.Ala1250Thr; Marsh et al., 2017; Marsh et al., 2018; Figure 8A and S6C). We further included two artificial mutant receptors that were previously shown to perturb NTN1 binding and chemoattraction (p.Val848Arg, p.His857Ala; Finci et al., 2014). First, these mutants were transfected into HEK293T and COS-7 cells that do not endogenously express DCC (Chen et al., 2013; Shekarabi and Kennedy, 2002). Immunoblotting and immunohistochemistry performed without cell permeabilisation revealed that all mutant DCC receptors were appropriately expressed and localised to the cell membrane (Gad et al., 2000; Figure S7E). Using a previously established *in vitro* binding assay (Müller and Soares, 2006; Zelina et al., 2014), we discovered that DCC mutant proteins with altered residues located at the NTN1 binding interface (p.V793G and p.G805E) were unable to bind NTN1 (Figure 8B), while all other receptors with altered residues lying outside of the NTN1 binding interface still bound NTN1 (p.M743L, p.V754M, p.A893T and p.M1217;A1250T; Figure 8B). Surprisingly, all eight mutant *DCC* receptors were unable to modulate cell morphology in the presence of NTN1 (Figure 8C-E; Table S1). Collectively, our results suggest a model whereby mutations that affect the ability for DCC to regulate cell shape (Figure 8F), are likely to cause callosal agenesis through perturbed MZG migration and IHF remodelling.

**Figure 8:**
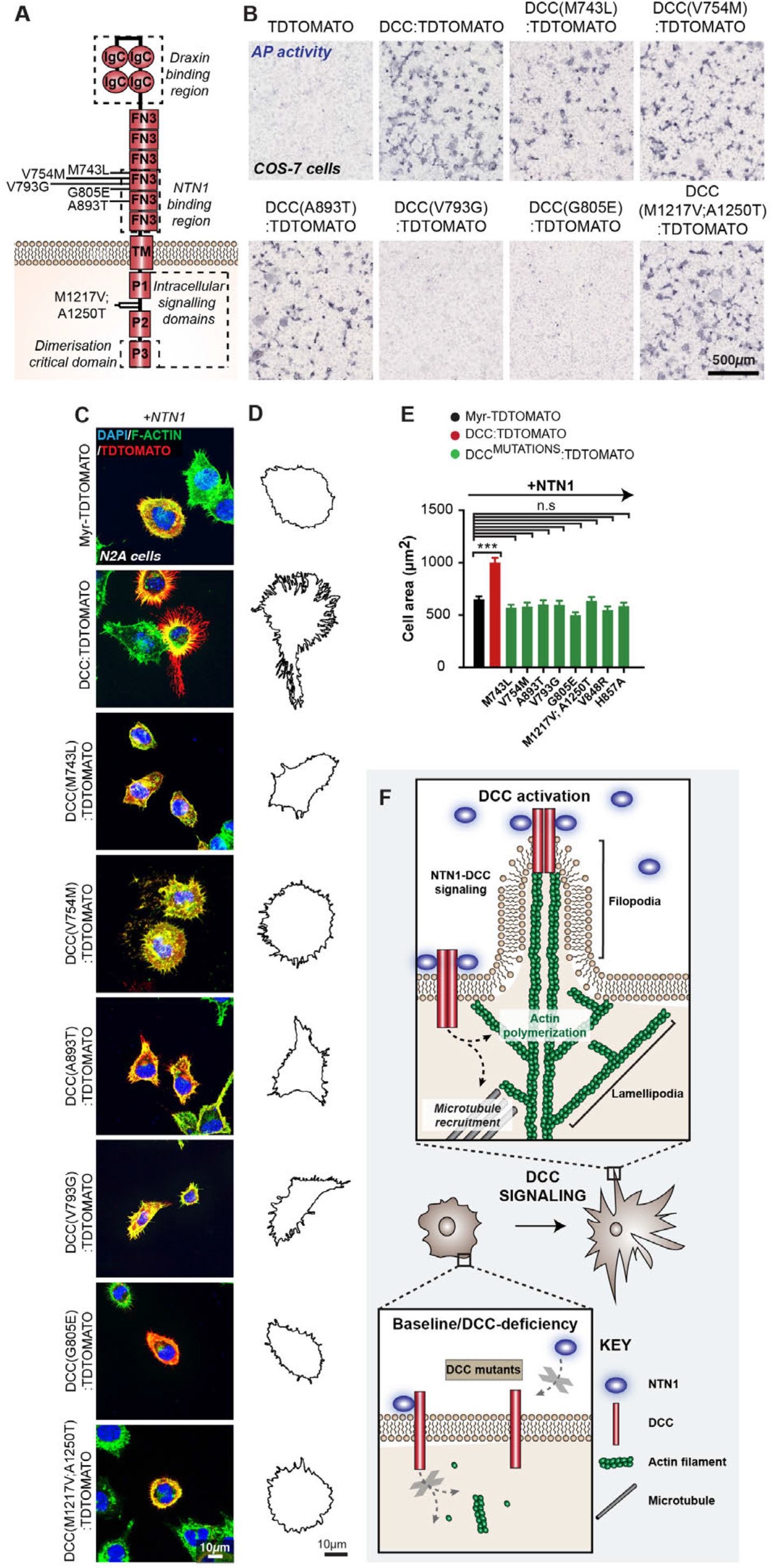
*DCC* mutations associated with human callosal agenesis are unable to modulate cell shape and show varied NTN1 binding. (A) Schema of transmembrane receptor DCC and its structural domains. Lines indicate the position of altered residues from missense *DCC* pathogenic variants identified in human individuals with CC abnormalities. FN3 = fibronectin type III-like domain, IgC = immunoglobulin-like type C domain, TM = transmembrane domain, P = P motif. (B) Colourimetric detection of alkaline phosphatase activity in COS-7 cells transfected with plasmids encoding TDTOMATO, DCC:TDTOMATO, and mutant DCC:TDTOMATO constructs, and incubated with a NTN1 alkaline phosphatase fusion protein. (C) Representative images of N2A cells immunolabelled for TDTOMATO (red), and F-actin (green) following transfection with plasmids encoding Myr-TDTOMATO, DCC:TDTOMATO, or DCC:TDTOMATO carrying missense mutations and stimulated with recombinant mouse NTN1 protein. (D) Outline of cell perimeter generated from images in B. (E) Quantification of the area of cells represented in B. Graph represents mean ± SEM. Statistics by Kruskal-Wallis test for multiple comparisons: n.s = not significant, ***p < 0.001. (F) Schema of model for DCC-mediated changes in cell shape: Activation of DCC by NTN1 induces dimerisation of the receptor and recruits intracellular signaling effectors to regulate actin polymerisation for filopodia and lamellipodia formation, and to regulate microtubule dynamics to promote membrane protrusions. Mutations that affect DCC signalling prevent DCC-mediated changes in cell shape. See related Figure S6 and Table S1.

## Discussion

Genes that encode axon guidance molecules frequently cause callosal dysgenesis when knocked out in mice (Edwards et al., 2014). This has led to the prevalent view that callosal dysgenesis in these mice might be primarily to due defects in callosal axon guidance towards and across the midline. Here, we identified a novel function for the classical axon guidance genes NTN1 and DCC in regulating the morphology of midline astroglia for IHF remodelling prior to CC and HC formation. Importantly, normal astroglial development and IHF remodelling are critical processes that precede and are necessary for subsequent CC axon guidance across the interhemispheric midline (Gobius et al., 2016). We find that defects in IHF remodelling are consistently associated with dysgenesis of the CC and HC in mice and humans with pathogenic variants in *Ntn1* or *Dcc*.

Our *in vitro* assays and analysis of mouse and human cell morphology indicate that the cytoskeletal remodelling function of NTN1-DCC signalling is likely to be crucial for MZG development, IHF remodelling, and subsequent CC formation. The timely differentiation and appropriate distribution of MZG cells at the IHF surface is required for their intercalation and IHF remodelling function (Gobius et al., 2016). Our data suggest a model whereby failed IHF remodelling associated with mutations in *Ntn1* and *Dcc* occurs due to delayed astroglial migration to the IHF as a consequence of perturbed process extension and organisation of MZG precursors. Notably, no medial extension of MZG processes across the basement membrane or perforations in the IHF to allow glia from each hemisphere to interact and intercalate were observed in *Ntn1* or *Dcc* mutant mice at any developmental stage examined. This suggests that NTN1-DCC signalling might also be required for MZG intercalation and removal of the intervening leptomeninges. The DCC homologue UNC-40 is known to facilitate formation of a polarised actin-rich cell protrusion in the *Caenorhabditis elegans* anchor cell, which breaches the basement membrane rich in UNC-6 (NTN1 homolog), enabling the cell to invade the vulval epithelium (Hagedorn et al., 2013; Ziel et al., 2009). DCC may perform a similar function in MZG by engaging secreted NTN1, which we found to be localised at the IHF basement membrane, by preferentially inducing polarisation of actin remodelling and process extension during MZG intercalation. Moreover, callosal axons that also rely on DCC-mediated cytoskeletal remodelling for growth and guidance, may non-cell-autonomously influence the final stages of MZG development via a secreted cue or contact-dependent mechanism. Further dissecting this would ideally involve even greater precision in complete and cell-type specific knockout of DCC and NTN1, since knockdown in a subset of cells or merely lowering the expression level was insufficient to induce a consistent phenotype.

Notably, we find that the P3 domain-dependent functions of DCC may be required for astroglial development and IHF remodelling. These functions include receptor dimerisation, interaction with the co-receptor ROBO1, or interaction with effectors FAK, MYO10, and TUBB3 (Fothergill et al., 2014; Li et al., 2004; Qu et al., 2013; Stein and Tessier-Lavigne, 2001; Wei et al., 2011; Xu et al., 2018). Accordingly, mice deficient in *Robo1, Fak* and *Tubb3*, as well as their signaling effectors *Cdc42, Fyn*, *Enah* and *Mena,* which normally act downstream of DCC to regulate the cell cytoskeleton, all display dysgenesis of the CC (Andrews et al., 2006; Beggs et al., 2003; Goto et al., 2008; Menzies et al., 2004; Tischfield et al., 2010; Yokota et al., 2010). Similarly, astroglial cells remodel their cytoskeleton to transition from a bipolar to multipolar morphology, and this process is known to involve the intracellular DCC effectors CDC42, RAC1, RHOA, N-WASP and EZRIN (Abe and Misawa, 2003; Antoine-Bertrand et al., 2011; Derouiche and Frotscher, 2001; Lavialle et al., 2011; Murk et al., 2013; Racchetti et al., 2012; Shekarabi et al., 2005; Zeug et al., 2018). Whether these molecules serve as downstream effectors of DCC to influence astroglial development and IHF remodelling during CC formation is an interesting question for future research.

In addition to NTN1 and DCC, as shown here, mice lacking the axon guidance molecules ENAH, SLIT2, SLIT3, and RTN4R have previously been reported to have incomplete IHF remodelling and disrupted midline glial development associated with callosal dysgenesis (Menzies et al., 2004; Unni et al., 2012; Yoo et al., 2017). Taken together, those studies and ours suggest that other axon guidance genes may play similar roles in astroglial development and IHF remodelling during CC formation. Additional candidate axon guidance molecules that may regulate IHF remodelling include EPHB1, EFNB3, GAP43, HS6ST1, HS2ST1, ROBO1 and VASP, since mouse mutants lacking these molecules display disrupted midline glial development and callosal dysgenesis (Andrews et al., 2006; Conway et al., 2011; Mendes et al., 2006; Menzies et al., 2004; Shen et al., 2004; Unni et al., 2012). Additional molecules of interest are EFNB1, EFNB3, EPHB2, and EPHA4, since these are expressed by MZG (Mendes et al., 2006).

In summary we have demonstrated that rather than solely regulating axon guidance during telencephalic commissure formation, *Dcc* and *Ntn1* are critical genes required for IHF remodelling. Moreover, our study provides a novel role for axon guidance receptor DCC in regulating astroglial morphology, organisation and migration. Exemplified by *Ntn1* and *Dcc,* our study provides support for widespread consideration of astroglial development and IHF remodelling as possible underlying mechanisms regulated by these and other classically regarded “axon guidance genes” during CC formation.

## Materials and Methods

### EXPERIMENTAL MODELS AND SUBJECT DETAILS

#### Animals

*Dcc^flox/flox^* (Krimpenfort et al., 2012)*, Dcc* knockout (Fazeli et al., 1997), *Dcc^kanga^* (Finger et al., 2002), *Emx1^iC**re**^* (Kessaris et al., 2006), *Ntn1-lacZ* (Serafini et al., 1996, and *tdTomato^flox_stop^* (Madisen et al., 2010) mice on the C57BL/6J background and CD1 wildtype mice were bred at The University of Queensland. Prior approval for all breeding and experiments were obtained from the University of Queensland Animal Ethics Committee. Male and female mice were placed together overnight and the following morning was designated as E0 if a vaginal plug was detected. *Dcc* knockout and *Dcc^kanga^* mice were genotyped by PCR and *Ntn1-lacZ* mice were tested for the presence of the *LacZ* gene and deemed homozygous if the β-galactosidase enzyme was trapped intracellularly, as previously described (Fazeli et al., 1997; Finger et al., 2002; Fothergill et al., 2014; Krimpenfort et al., 2012; Serafini et al., 1996). *Dcc^flox/flox^* mice were genotyped by the Australian Equine Genetics Research Centre at the University of Queensland.

#### Human subjects

Ethics for human experimentation was acquired by local ethics committees at The University of Queensland (Australia), the Royal Children’s hospital (Australia), and UCSF Benioff Children’s Hospital (USA). Genetic studies were performed previously (Marsh et al., 2017). Structural MR images were acquired as previously described (Marsh et al., 2017). In our study, we analysed the brain phenotype of affected individuals in family 2 (carrying *DCC* p.Val793Gly) and family 9 (carrying *DCC* p.Met1217Val;p.Ala1250Thr in cis) from our previous study.

### METHOD DETAILS

#### Cell birth-dating and tissue collection

For cell birth dating studies, 5-ethynyl-2’-deoxyuridine (EdU; 5 mg per kg body weight, Invitrogen) dissolved in sterile phosphate buffer solution (PBS) was injected into the intraperitoneal cavity of awake pregnant dams. Brains were fixed via transcardial perfusion or immersion fixation with 4% paraformaldehyde (PFA).

#### Cell culture

All cell lines were cultured at 37°C within a humidified atmosphere containing 5% CO_2_ and immersed in Dulbecco’s Modified Eagles Medium (DMEM) medium (Invitrogen or HyClone™), supplemented with 10% fetal bovine serum. U251 cells were plated on poly-d-lysine-coated coverslips (via submersion in 0.05 mg/mL solution, Sigma-Aldrich) at 10% confluence 24 hours prior to transfection. The pCAG-TDTOMATO, pCAG-H2B-GFP-2A-MyrTDTOMATO, pCAG-DCC:TDTOMATO and pCAG-DCC^kanga^:TDTOMATO plasmids (1 µg) were transfected into the plated U251 cells using FuGENE 6 (Promega) in Opti-MEM (Gibco, Life Technologies). Cells were then grown for 20 hours and either fixed with 4% paraformaldehyde/4% sucrose or stimulated with ligand. Since 100ng/mL of recombinant NTN1 is sufficient to induce morphological changes in primary oligodendrocyte precursor cells (Rajasekharan et al., 2009), 200ng of recombinant mouse NTN1 protein (R&D Systems) was diluted in sterile PBS and added to cultures within 2 mL media. When ligand was added, cells were grown for a further 12 hours before fixation with 4% PFA/4% sucrose. N2A cells were cultured and transfected as outlined for the U251 cells except that after NTN1 stimulation, cells were cultured for only 8 hours before fixation. The pCAG-DCC:TDTOMATO wildtype and missense mutant receptor constructs (1.764 µg) were also transfected into HEK293T cells cultured on acid-washed coverslips using FuGENE HD (Promega) in Optimem (Gibco, Life Technologies). After 24 hours, cells were fixed with 4% PFA/4% sucrose.

#### NTN1-binding assay

Supernatant containing alkaline phosphatase-conjugated *NTN1* (*NTN1-AP*) was generated from expression in HEK293T cells as previously described (Zelina et al., 2014). The pCAG-DCC:TDTOMATO wildtype and missense mutant receptor constructs (0.2 µg) were transfected into COS-7 cells using Lipofectamine® 2000 (Invitrogen). After 48 hours, cells were incubated with *NTN1*-AP supernatant (1:50) for 90 minutes at room temperature. Cells were washed and *NTN1*-binding activity was determined using colorimetric detection as previously described (Zelina et al., 2014).

#### Western blot

The pCAG-DCC:TDTOMATO wildtype and missense mutant receptor constructs (0.2 µg) were transfected into COS-7 cells using Lipofectamine® 2000 (Invitrogen). After 48 hours, the cells were lysed and proteins separated using gel electrophoresis. A western blot was performed to detect DCC expression levels using a goat polyclonal anti-DCC antibody (1:200, ab 18207, Santa Cruz Biotechnology). Gapdh was used as a loading control and was detected using a rabbit monoclonal anti-Gapdh antibody (1:2000, Cell Signaling Technology).

#### Immunohistochemistry

Brain sections were processed for standard fluorescence immunohistochemistry as previously described (Moldrich et al., 2010) with the following minor modifications: All sections were post-fixed on slides with 4% PFA and then subjected to antigen retrieval (125°C for 4 minutes at 15 psi in sodium citrate buffer) prior to incubation with primary antibodies. Alexa Fluor IgG (Invitrogen), horseradish peroxidase-conjugated (Millipore) or biotinylated (Jackson Laboratories) secondary antibodies, used in conjunction with Alexa Fluor 647-conjugated Streptavidin (Invitrogen) amplification were used according to the manufacturer’s instructions. EdU labeling was performed using the Click-iT EdU Alexa Fluor 488 Imaging Kit (Invitrogen). Cell nuclei were labeled using 4’,6-diamidino-2-phenylindole, dihydrochloride (DAPI, Invitrogen) and coverslipped using ProLong Gold anti-fade reagent (Invitrogen) as mounting media. Primary antibodies used for immunohistochemistry were: rabbit anti-APC (1:250, ab15270, Abcam), mouse anti-α-dystroglycan (1:250, clone IIH6C4, 05-593, Merk), rabbit anti-β-catenin (1:500, 9562, Cell signaling technology), mouse anti-β-dystroglycan (1:50, MANDAG2, 7D11, Developmental studies hybridoma bank), chicken anti β-galactosidase (1:500, ab9361, Abcam), rabbit anti-cleaved-caspase3 (1:500, 9661, cell signaling technology), donkey anti-DCC (1:500, ab 18207, Santa cruz biotechnology), mouse anti-Gap43 (1:500; AB1987, Millipore), mouse anti-Gfap (1:500; MAB3402, Millipore), rabbit anti-Gfap (1:500; Z0334, Dako), mouse anti-Glast (or Eaat1; 1:500; ab49643, Abcam), rabbit anti-Glast (or Eaat1; 1:250; ab416, Abcam), mouse anti-Ki67 (1:500; 550609, BD Pharmingen), chicken anti-Laminin (1:500; LS-C96142, LSBio), rabbit anti-Laminin (1:500; L9393, Sigma), mouse anti-N-cadherin (1:250, ab610921, BD Biosciences), rat anti-Nestin (AB 2235915, DSHB), chicken anti-Nestin (1:1000, ab134017, Abcam), goat anti-Netrin1 (AF1109, R&D Systems), mouse anti-Neurofilament (1:500; MAB1621, Chemicon), rabbit anti-Nfia (1:500; ARP32714, Aviva Systems Biology), rabbit anti-Nfib (1:500; HPA003956, Sigma), rabbit anti-neuronal-specific-ßIII-tubulin (1:500; ab18207, Abcam), rabbit anti-phospho p44/42 Mapk (or Erk1/2; 1:250; 9101, Cell Signaling), rabbit anti-Sox9 (1:500, AB553, Merck), and goat anti-TDTOMATO (1:500, ab8181-200, Sicgen). For actin staining, Alexa fluor-conjugated phalloidin (A22287, Thermofisher scientific) was incubated on tissue for thirty minutes in the dark as per the manufacturer’s instructions, prior to the addition of primary antibodies. Immunohistochemistry was performed in a similar manner for cultured cells, with the following minor exceptions: HEK293T cells expressing wildtype and mutant pCAG-DCC:TDTOMATO constructs were not permeabilized to confirm exogenous DCC receptor localisation to the plasma membrane.

#### In situ hybridization

In situ hybridization was performed as previously described (Moldrich et al., 2010), with the following minor modifications: Fast red (Roche) was applied to detect probes with fluorescence. The *Fgf8* cDNA plasmid was a kind gift from Gail Martin, University of California, San Francisco. The *Ntn1* cDNA plasmid was provided by the Cooper lab. The *Mmp2* cDNA plasmid was generated by the Richards lab with the following primers: forward 5’ - GAAGTATGGATTCTGTCCCGAG – 3’ and reverse 5’ – GCATCTACTTGCTGGACATCAG – 3’. The *Dcc* cDNA plasmid was generated by the Richards lab with the following primers, courtesy of the Allen Developing Brain Atlas: forward 5’ - ATGGTGACCAAGAACAGAAGGT - 3’ and reverse 5’ – AATCACTGCTACAATCACCACG – 3’.

#### Plasmid expression constructs for cell culture and in utero electroporation

A TDTOMATO fluorophore (Clontech) was subcloned into a *pCAG* backbone to generate the pCAG-TDTOMATO plasmid. pCAG-H2B-GFP-2A-MyrTDTOMATO was provided by Arnold Kriegstein (University of California San Francisco). The *Dcc*-shRNA construct was provided by Xiong Zhiqi (Chinese Academy of Sciences, Shanghai; shRNA 1355 in Zhang et al., 2018). The *Dcc*-CRISPR nickase constructs were designed using the ATUM tool and obtained from ATUM to target *Dcc* exon 2 (*Dcc*-CRISPR 1, targeting chr18:71,954,969 - 71,955,009) and *Dcc* exon 3 (*Dcc*-CRISPR 2, targeting chr18:71,826,146 - 71,826,092). *Dcc*-CRISPR 1 had the maximum target score across the whole DCC coding sequence, while *Dcc*-CRISPR 1 had the maximum target score within exon 3 only.

To generate the pCAG-DCC:tdTomato plasmid, DCC:TDTOMATO (pmDCC:TDTOMATO; provided by Erik Dent, University of Wisconsin-Madison) was subcloned into the pCag-DsRed2 plasmid (Addgene, 15777, Cambridge, MA), by excising DsRed2.

For site-directed mutagenesis, the QuickChange II Site-Directed Mutagenesis Kit (Stratagene, Catalogue #200524) was used in accordance with the manufacturer’s instructions. The following primer pairs were used for site-directed mutagenesis: p.Met743Leu: Forward 5’- GAGGAGGTGTCCAACTCAAGATGATACAGTTTGTCTG – 3’, reverse 5’ – CAGACAAACTGTATCATCTTGAGTTGGACACCTCCTC – 3’. p.Val754Met: Forward 5’ – TAATATAGCCTCTCACCATGATGTTTGGGTTGAGAGG – 3’, reverse 5’ – CCTCTCAACCCAAACATCATGGTGAGAGGCTATATTA – 3’. p.Ala893Thr: Forward 5’ – ACTTGTACTTGGTACTGGCAGAAAAGCTGGTCCT – 3’, reverse 5’ – AGGACCAGCTTTTCTGCCAGTACCAAGTACAAGT – 3’. p.Val793Gl: Forward 5’ – ACTAGAGTCGAGTTCTCATTATGGAATCTCCTTAAAAGCTTTCAAC -3’, reverse 5’ –GTTGAAAGCTTTTAAGGAGATTCCATAATGAGAACTCGACTCTAGT – 3’. p.Gly805Glu: Forward 5’ – CACTTTCGTAGAGAGGGACCTCTTCTCCGGCATTGTTGAA – 3’, reverse 5’ – TTCAACAATGCCGGAGAAGAGGTCCCTCTCTACGAAAGTG – 3’. p.Met1217Val;p.Ala1250Thr: Forward 1 5’ – GTTCCAAAGTGGACACGGAGCTGCCTGCGTC – 3’, reverse 1 5’ – GACGCAGGCAGCTCCGTGTCCACTTTGGAAC – 3’, forward 2 5’ – GTACAGGGATGGTACTCACAACAGCAGGATTACTGG – 3’, reverse 2 5’ - CCAGTAATCCTGCTGTTGTGAGTACCATCCCTGTAC – 3’. p.Val848Arg: Forward 5’ – CAGCCTGTACACCTCTTGGTGGGAGCATGGGGG – 3’, reverse 5’ – CCCCCATGCTCCCACCAAGAGGTGTACAGGCTG – 3’. p.His857Ala: Forward 5’ – ACCCTCACAGCCTCAGCGGTAAGAGCCACAGC – 3’, reverse 5’ – GCTGTGGCTCTTACCGCTGAGGCTGTGAGGG- 3’. p.del-P3(Kanga): Forward 5’ – CCACAGAGGATCCAGCCAGTGGAGATCCACC – 3’, reverse 5’ – GGTGGATCTCCACTGGCTGGATCCTCTGTGG – 3’.

#### In utero electroporation

In utero electroporation was performed as previously described (Suárez et al., 2014). Briefly, 2 µg/µL of *Dcc*-shRNA or *Dcc*-CRISPR were combined with 0.5 µg/µL TDTOMATO and 0.0025% Fast Green dye, and then microinjected into the lateral ventricles of E13 *Dcc^kanga^* embryos. 5, 35 V square wave pulses separated by 100 ms were administered with 3mm paddles over the head of the embryo to direct the DNA into the cingulate cortex. Embryos were collected at E18 for analysis.

#### Image acquisition

Confocal images were acquired as either single 0.4-0.9 µm optical sections or multiple image projections of ∼15-20 µm thick z-stacks using either an inverted Zeiss Axio-Observer fitted with a W1 Yokogawa spinning disk module, Hamamatsu Flash4.0 sCMOS camera and Slidebook 6 software, or an inverted Nikon TiE fitted with a Spectral Applied Research Diskovery spinning disk module, Hamamatsu Flash4.0 sCMOS camera and Nikon NIS software. Alternatively, for images of HEK293T cells, a LSM 780 confocal microscope was used. For imaging of NTN1-AP binding, a NanoZoomer 2.0-HT whole slide imager was used in conjunction with Hamamatsu (NDP_Viewer) software. Images were pseudocolored to permit overlay, cropped, sized, and contrast-brightness enhanced for presentation with ImageJ and Adobe Photoshop software.

#### Measurements and cell quantification

Measurements of IHF length were performed using ImageJ v1.51s freeware (National Institutes of Health, Bethsda, USA). The length of the IHF within the interhemispheric midline was determined by comparing Laminin and DAPI-staining. To account for inter-brain variability, this length was then normalised to the entire length of the telencephalon along the interhemispheric midline, which was measured from the caudal-most point of the telencephalon to the rostral edge of cerebral hemispheres.

Cell proliferation and cell death in *Dcc^kanga^* MZG was automatically counted using Imaris software (Bitplane) from a region of interest delineated by Glast staining, and excluding the IHF within a single z-slice.

The number of Sox9-positive cell bodies was counted manually using the Cell Counter plugin in ImageJ v1.51s freeware. Cell proliferation and cell death in tissue was automatically counted using Imaris software (Bitplane) from a region of interest delineated by Glast staining that excluded the IHF in a single z-slice.

The perimeter, circularity and area of U251 and N2A cells was measured from mean intensity projections of TDTOMATO images following thresholding in ImageJ v1.51s freeware (National Institutes of Health, Bethsda, USA). 48-191 cells per condition were analysed from 3-5 biological replicates.

#### Fluorescence intensity analysis

To compare fluorescence intensity, tissue sections were processed under identical conditions for immunofluorescence. Fluorescent images at 20x or 40x magnification were acquired using identical exposure settings for each fluorescent signal and identical number of slices through the z plane. A multiple intensity projection was created for each z-stack to create a 2D image. Identical regions of interest were outlined in ImageJ freeware and the fluorescence intensity was plotted versus the distance.

#### Quantification and statistical analysis

A minimum of three animals were analysed for each separate phenotypic analysis. Sex was not determined for embryonic studies. A mix of male and female adult mice were used to determine the length of the IHF in *Dcc^kanga^* and C57Bl/6 mice. All measurements and cell counting were performed on deidentified files so the researcher remained blind to the experimental conditions. For comparison between two groups, the data was first assessed for normality with a D’Agostino-Pearson omnibus normality test and then statistical differences between two groups were determined either with a parametric Student’s t-test, or a non-parametric Mann-Whitney test in Prism software (v.6-v.8; GraphPad). For multiple comparisons of cell culture conditions, a Kruskal-Wallis test was performed with post-hoc Dunn’s multiple comparison test. p ≤ 0.05 was considered significantly different, where all p values are reported in text. All values are presented as mean ± standard error of the mean (SEM).

### CONTACT FOR REAGENT AND RESOURCE SHARING

Further information and requests for resources and reagents should be directed to and will be fulfilled by Professor Linda J Richards.

### DATA AVAILABILITY

Microscopy data, measurements and statistical analyses are available. This study did not generate code.

## Acknowledgements

We thank Marc Tessier-Lavigne for the *Ntn1-lacZ*, and *Dcc* knockout mouse lines, and Susan Ackermann (Jackson Laboratory) for the *Dcc^kanga^* mouse lines. We thank colleagues for providing the constructs listed. We thank Luke Hammond, Rumelo Amor, Arnaud Guardin and Andrew Thompson for assistance with microscopy, which was performed in the Queensland Brain Institute’s Advanced Microscopy Facility. This work was supported by Australian NHMRC grants GNT456027, GNT631466, GNT1048849 and GNT1126153 to L.J.R and GNT1059666 to P.J.L and R.J.L, and USA National Institutes of Health grant 5R01NS058721 to E.S and L.J.R. R.S received an Australian Research Council DECRA fellowship (DE160101394). A.P.L.M and L.M were supported by a Research training program scholarship (Australian Postgraduate Award). A.P.L.M was further supported by an NHMRC Early Career Research Fellowship (APP1156820). AL.S.D was supported by an Australian Postgraduate award. L.M and AL.S.D received a Queensland Brain Institute Top-Up Scholarship. L.R.F was supported by a UQ Development Fellowship, R.J.L. was supported by a Melbourne Children’s Clinician Scientist Fellowship and L.J.R. was supported by an NHMRC Principal Research Fellowship (GNT1120615).

We thank the families and members of the Australian Disorders of the Corpus Callosum (AusDoCC) for their support and time in being involved in this research. We thank the International Research Consortium for the Corpus Callosum and Cerebral Connectivity (IRC5, https://www.irc5.org) researchers for discussions and input.

## Author contributions

Conceptualization, L.M, I.G, A.P.L.M, R.S, L.R.F, H.M.C, E.S, P.J.L, R.J.L, and L.J.R; Investigation, L.M, I.G, A.P.L.M, R.S, L.R.F, C.B., Y.Y, S.S, Y.Z., A.D, AL.D, P.K., T.F, T.J.E and L.J.R; Writing – original draft and visualization, L.M, I.G, A.P.L.M, and L.J.R; Writing – review and editing, all authors; Supervision, I.G, R.S, P.K, A.C,Y.Z, R.J.L, P.J.L and L.J.R. Funding acquisition, L.J.R, E.S, A.C, P.J.L, and R.J.L.

