## Supplementary figures and tables for "The DCC receptor regulates astroglial development essential for telencephalic morphogenesis and corpus callosum formation"

**Supplemental figures**

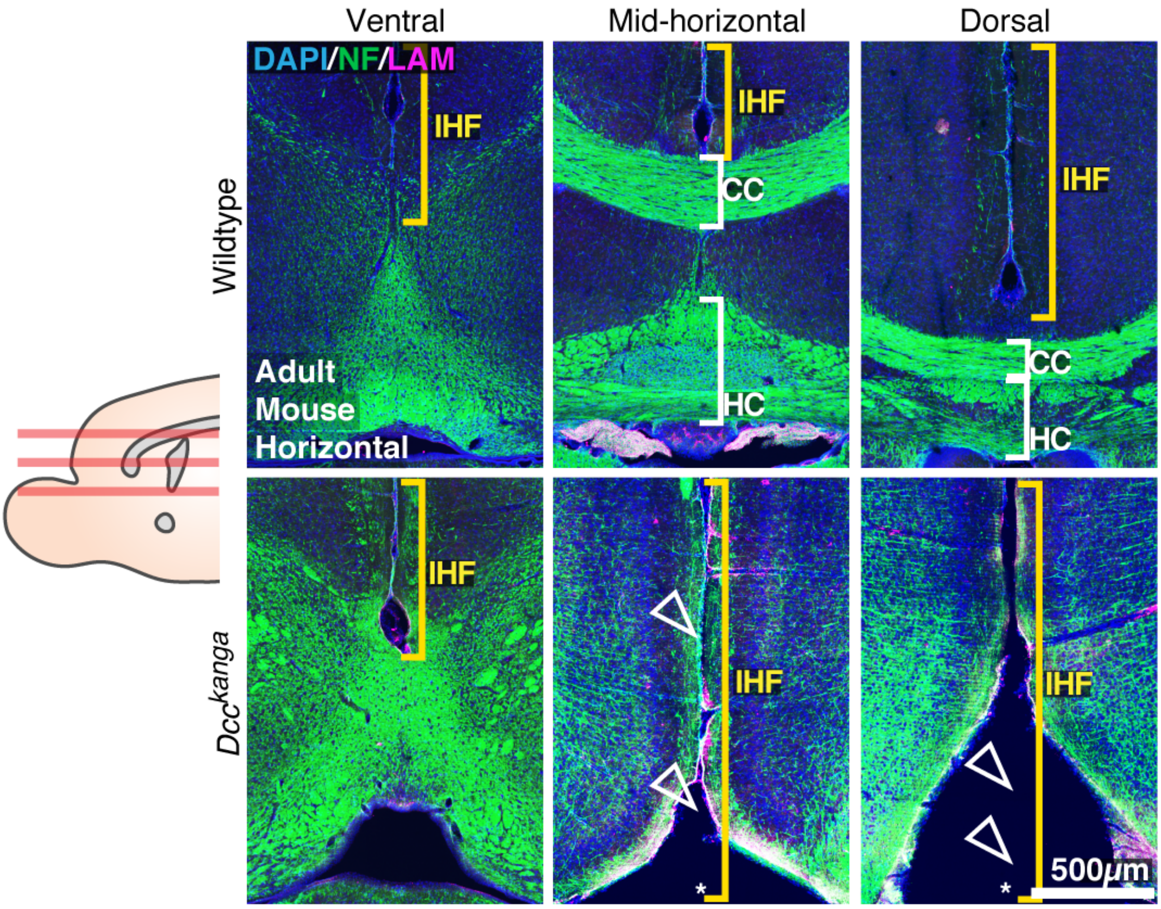


**Figure S1: The IHF is not remodelled in adult *Dcc*$^{kanga}$ mice**

Neurofilament (NF)-positive axons (green) and pan-Laminin (LAM)-positive leptomeninges and basement membrane (magenta) in adult wildtype and *Dcc*$^{kanga}$ mice reveal presence/absence of the CC and HC (white brackets and arrowheads), the extent of the IHF (yellow brackets) and absence of the septal substrate in *Dcc*$^{kanga}$ mice (asterisks).

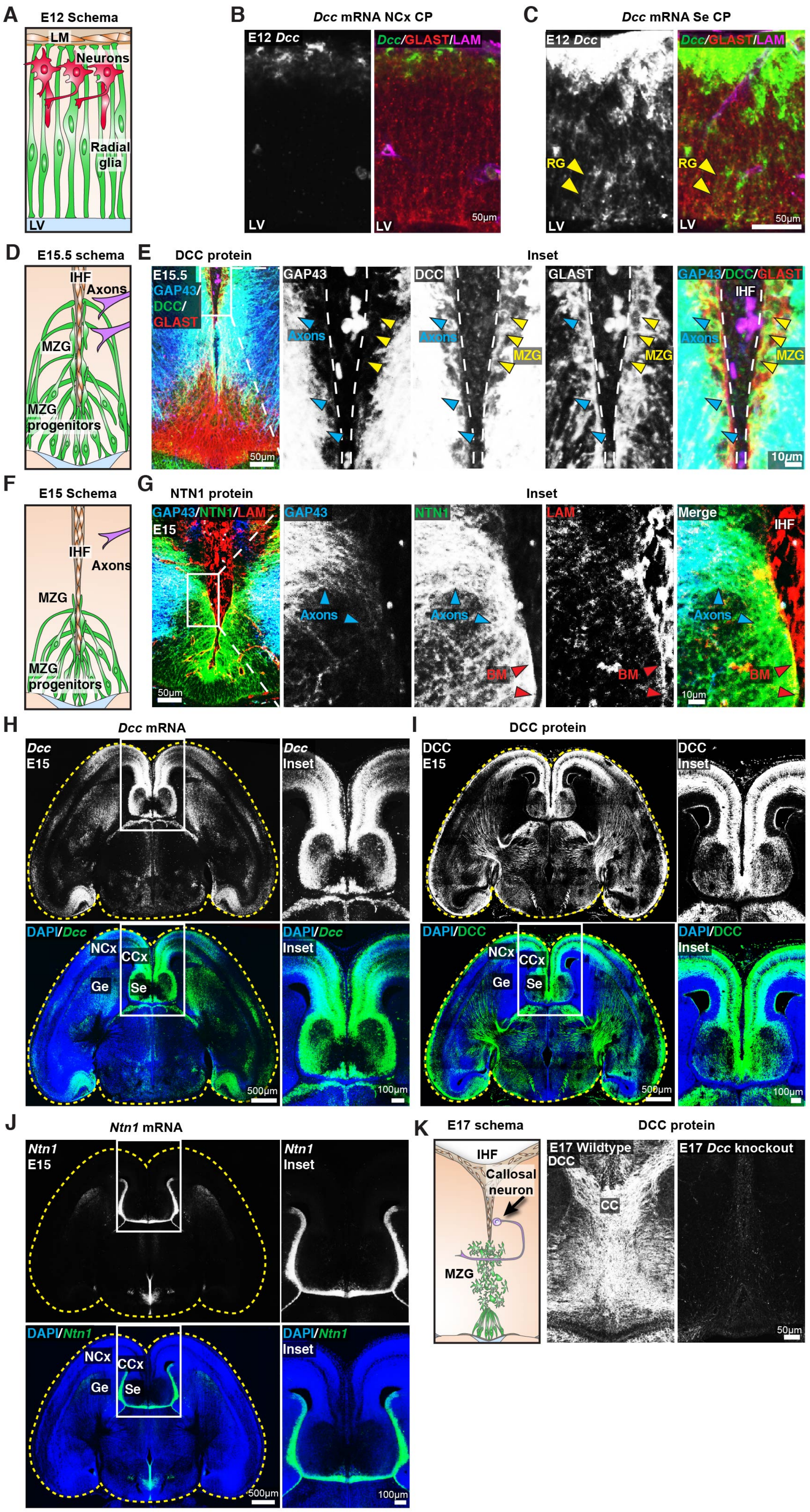

A
E12 Schema
LM
Neurons
Radial glia
LV
B
Dcc mRNA NCx CP
E12 Dcc
Dcc/GLAST/LAM
LV
50µm
C
Dcc mRNA Se CP
E12 Dcc
Dcc/GLAST/LAM
RG
LV
50µm
D
E15.5 schema
IHF
Axons
MZG
MZG progenitors
E
DCC protein
Inset
E15.5 GAP43/ DCC/ GLAST
GAP43
DCC
GLAST
GAP43/DCC/GLAST
IHF
Axons
MZG
50µm
10µm
F
E15 Schema
IHF
Axons
MZG
MZG progenitors
G
NTN1 protein
Inset
GAP43/NTN1/LAM
E15
GAP43
NTN1
LAM
Merge
IHF
Axons
BM
50µm
10µm
H
Dcc mRNA
Dcc E15
Dcc Inset
DAPI/Dcc
DAPI/Dcc Inset
NCx
CCx
Ge
Se
500µm
100µm
I
DCC protein
DCC E15
DCC Inset
DAPI/DCC
DAPI/DCC Inset
NCx
CCx
Ge
Se
500µm
100µm
J
Ntn1 mRNA
Ntn1 E15
Ntn1 Inset
DAPI/Ntn1
DAPI/Ntn1 Inset
NCx
CCx
Ge
Se
500µm
100µm
K
E17 schema
DCC protein
IHF
Callosal neuron
MZG
E17 Wildtype DCC
E17 Dcc knockout
CC
50µm

**Figure S2: DCC is expressed in MZG**

(A, D and F) Schemas of key cellular components within the telencephalic midline.

(B and C) *Dcc* mRNA (green), Glast-positive glia (red), and pan-laminin (LAM)-positive leptomeninges and basement membrane (magenta) across the cortical plate (Cp) within the neocortex (NCx) or septum (Se) in horizontal sections of wildtype mice reveals *Dcc*-positive/Glast-positive radial glial (RG) fibers (yellow arrowheads). LV = lateral ventricle.

(E) DCC protein (green), Gap43-positive axons (blue), and Glast-positive MZG (red) in horizontal sections of E15 wildtype mice (right panels), indicate DCC-positive/Glast-positive cells (yellow arrowheads) and DCC-positive/Gap43-positive axons (blue arrowheads) that are approaching the midline and are adjacent to MZG.

(G) Gap43-positive axons (blue), NTN1 protein (green), and pan-Laminin (LAM)-positive leptomeninges and basement membrane (magenta) in horizontal sections of E15 wildtype mice reveal NTN1-positive/Gap43-positive axons (blue arrowheads) approaching the midline and NTN1-positive/LAM-positive basement membrane (BM; red arrowheads) of the IHF.

(H, I and J) Mid-horizontal tissue sections encompassing the entire telencephalon (yellow outlines) with in situ hybridization for *Dcc* mRNA or *Ntn1* mRNA or immunohistochemistry for DCC protein (all white or green), counterstained with DAPI (blue). Insets of the telencephalic midline are shown on the right.

(K) DCC immunohistochemistry in horizontal sections of E17 wildtype and *Dcc* knockout mice with schema of key cellular components within the telencephalic midline.

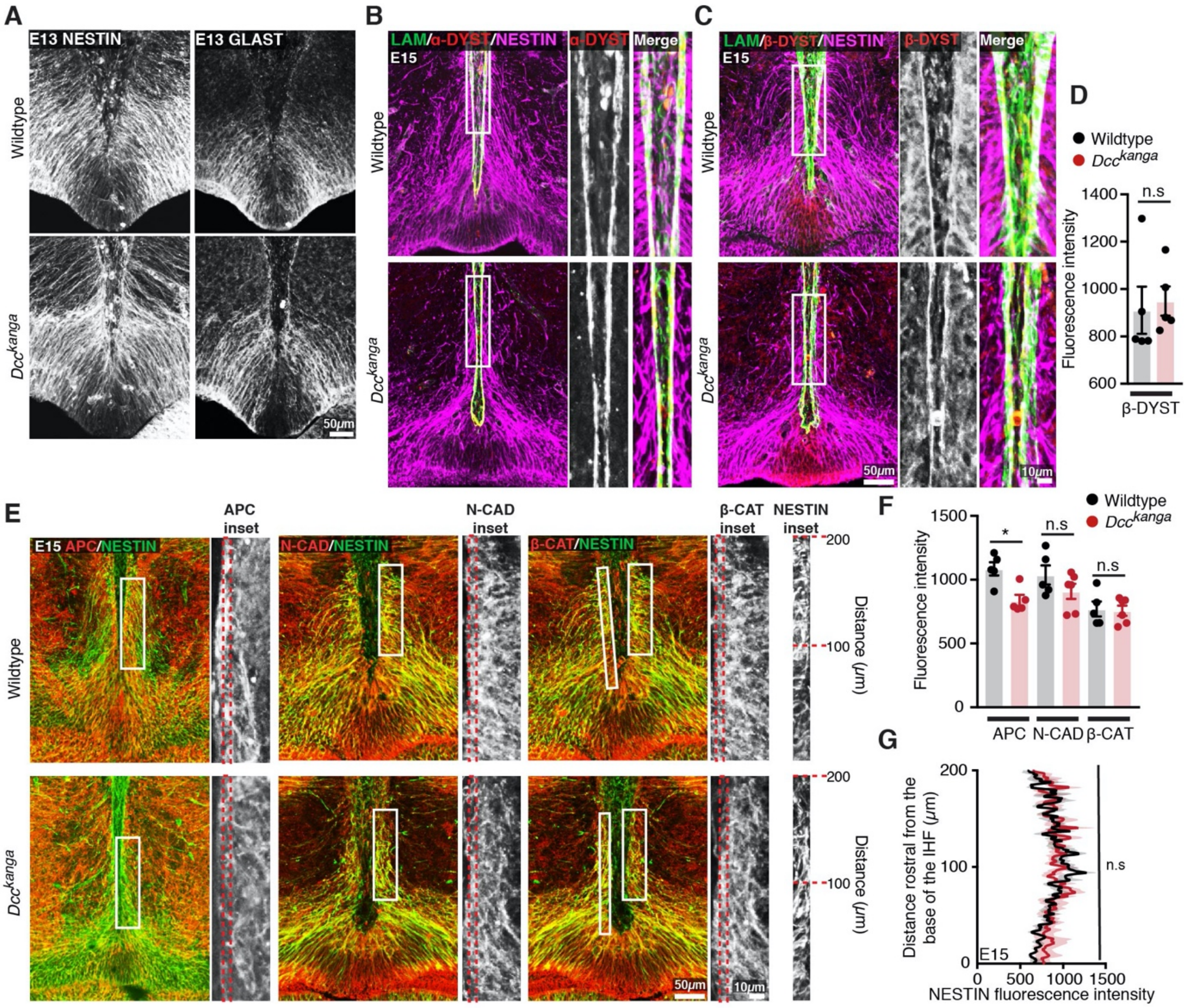


**Figure S3: DCC is not required for endfeet attachment or molecular polarity of MZG**

(A) Nestin-positive radial glia (white), and Glast-positive MZG (white) in horizontal sections of E13 wildtype and *Dcc*$^{kanga}$ mice.

(B and C) Pan-Laminin (LAM)-positive leptomeninges and basement membrane (green), Nestin-positive radial glia (magenta), and α-dystroglycan (α-DYST; red; B), or β-dystroglycan (β-DYST; red, C) in horizontal sections of E15 wildtype and *Dcc*$^{kanga}$ mice.

(D) Quantification of fluorescence intensity of β-DYST along 200 $\mu$m of the IHF surface as outlined with red dotted box in C.

(E) Nestin-positive radial glia (green) with either Adenomatous polyposis coli (APC, red), N-cadherin (N-CAD; red), or β-catenin (β-CAT; red) in horizontal sections of E15 wildtype and *Dcc*$^{kanga}$ mice with insets.

(F) Quantification of the fluorescence intensity of APC, N-CAD and β-CAT within 5 $\mu$m of the IHF as outlined in red dotted-edged boxes from E.

(G) Quantification of the fluorescence intensity of Nestin-positive radial glial endfeet within 5 $\mu$m of the IHF surface from inset in E.

All graphs represent mean ± SEM. Statistics by Mann-Whitney test: n.s = not significant, $^*p < 0.05$. See related Table S1.

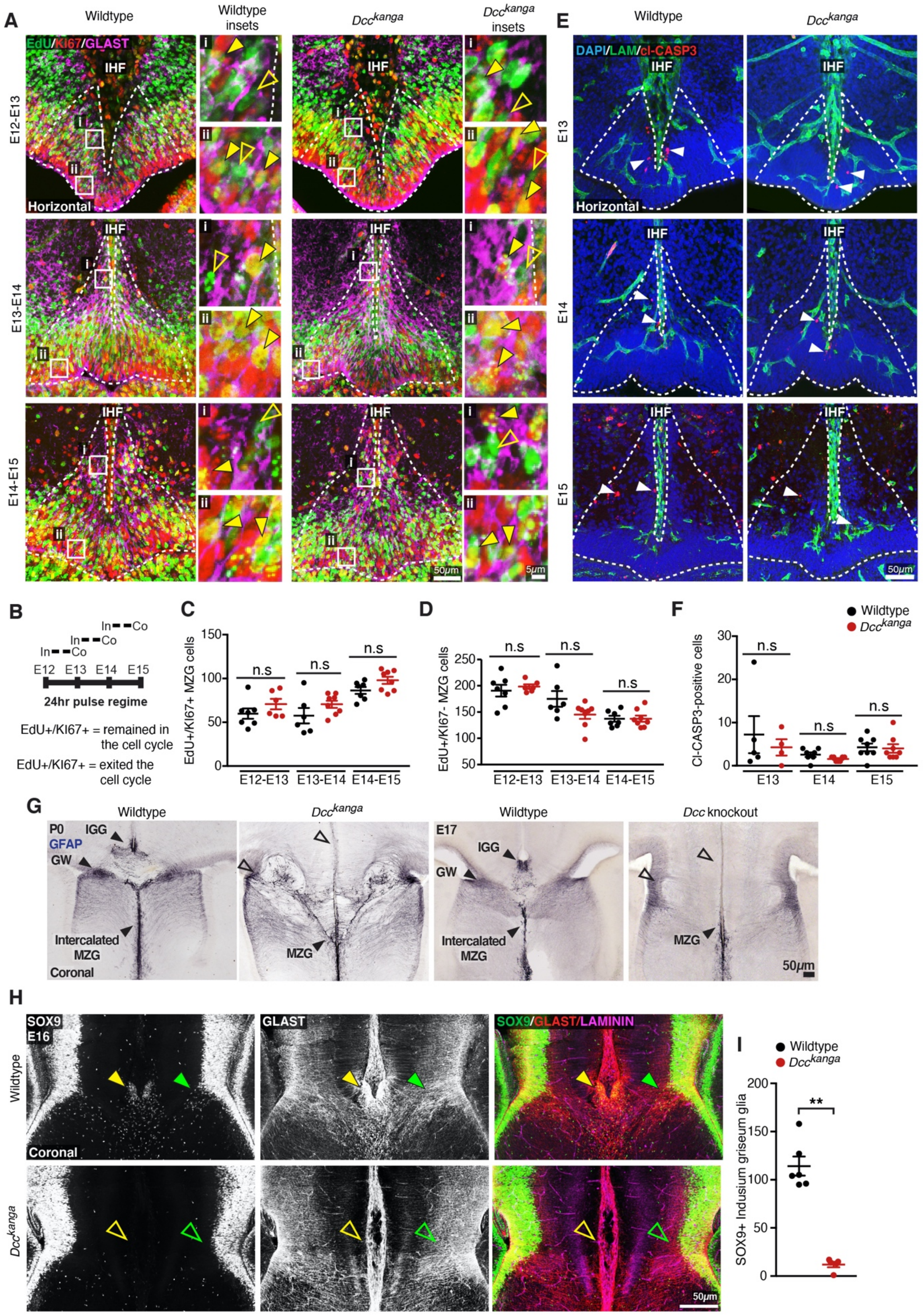
A
Wildtype
Wildtype insets
Dcc kanga
Dcc kanga insets
EdU/KI67/GLAST
IHF
Horizontal
E12-E13
E13-E14
E14-E15
i
ii
50μm
5μm
E
DAPI/LAM/cl-CASP3
E13
E14
E15
B
In Co
E12 E13 E14 E15
24hr pulse regime
EdU+/KI67+ = remained in the cell cycle
EdU+/KI67+ = exited the cell cycle
C
EdU+/KI67+ MZG cells
n.s
E12-E13 E13-E14 E14-E15
D
EdU+/KI67- MZG cells
F
Wildtype
Dcc kanga
Cl-CASP3-positive cells
E13 E14 E15
G
P0
GFAP
IGG
GW
Intercalated MZG
Coronal
MZG
E17
Dcc knockout
H
SOX9
E16
GLAST
SOX9/GLAST/LAMININ
I
SOX9+ Indusium griseum glia
**

**Figure S4: DCC does not regulate the proliferation or cell death of MZG but regulates the formation of the indusium griseum glia and glial wedge**

(A) Mouse MZG cells were birth-dated with the thymidine analog EdU every 24 hours, from E12 to E15 in wildtype and *Dcc*$^{kanga}$ mice. Representative images of EdU (green), cell cycle marker, Ki67 (red), and MZG marker Glast (magenta) are shown for wildtype and *Dcc*$^{kanga}$ mice, with the distribution of MZG progenitors within the telencephalic hinge niche outlined with white dotted lines. Yellow arrowheads in insets point out EdU cells that are either Ki57-positive (filled arrowheads) or Ki67-negative (open arrowheads) in selected insets. The number of cells expressing each marker is quantified in (C) and (D).

(B) Schema of the EdU injection (In) and collection (Co) regime and interpretation of co-labelled and non-co-labelled cells.

(E) Laminin(LAM)-positive leptomeninges and basement membrane (green) and cleaved-caspase3-positive apoptotic cells (red, white arrowheads) in E13-E15 wildtype and *Dcc*$^{kanga}$ mice. The number of cleaved-caspase3 (Cl-CASP3)-positive cells within the telencephalic hinge niche (white dotted lines) is quantified in (F).

(G) Mature astroglial marker GFAP in coronal sections of P0 *Dcc*$^{kanga}$ mice and E17 *Dcc* knockout mice and their wildtype littermates reveals midline glial populations, the glial wedge (GW), the indusium griseum glia (IGG), and the MZG (filled arrowheads) or their absence/malformation (open arrowheads).

(H) Glial-specific cell body marker SOX9 (white or green), glial cell membrane marker Glast (white or red), and IHF marker Laminin (magenta) in E16 coronal sections from *Dcc*$^{kanga}$ mice indicate the presence or absence of SOX9-positive/Glast-positive cell bodies at the pial surface of the IHF (yellow arrowheads) and within the intermediate zone (green arrowheads).

(I) Quantification of SOX9-positive IGG cell bodies at the pial surface of the IHF in E16 wildtype and *Dcc*$^{kanga}$ mice from immunohistochemistry in G.

All graphs represent mean ± SEM. Statistics by Mann-Whitney test: n.s = not significant, $^{**}p < 0.01$. See related Table S1.

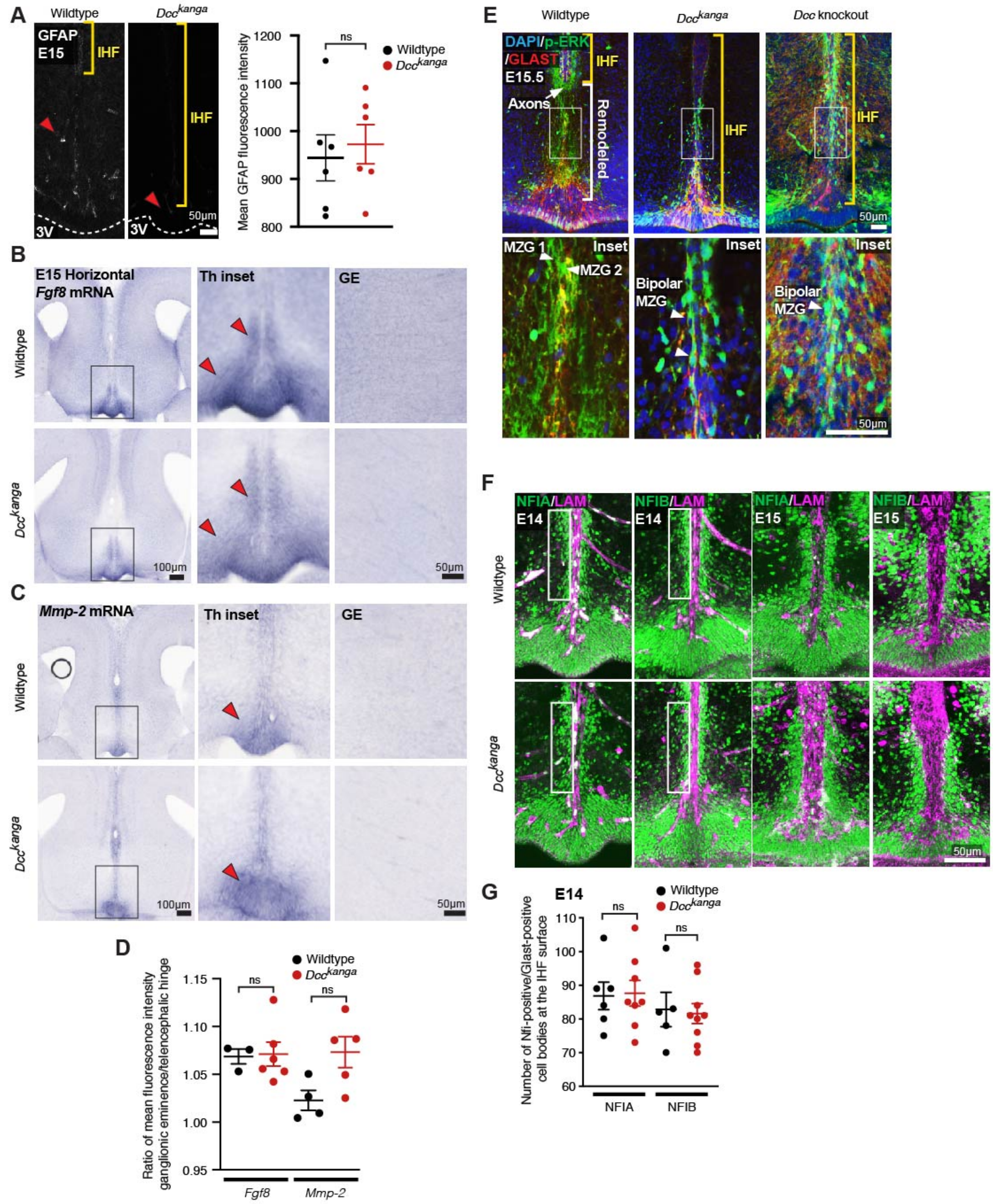

**Figure S5: DCC is not required for astroglial differentiation of MZG**

(A) Mature astroglial marker Gfap (white) in horizontal sections of E15 wildtype and *Dcc*$^{kanga}$ mice with quantification of Gfap average fluorescence intensity. The surface of the third ventricle (3V) is outlined with dotted white lines. Red arrowheads indicate reactive blood vessels that are not Gfap-positive glia. Yellow brackets indicate the position of the interhemispheric fissure (IHF).

*Fgf8* mRNA (B) or *Mmp-2* mRNA (C) in horizontal sections of E15 wildtype and *Dcc*$^{kanga}$ mice. Red arrowheads indicate reactivity in the telencephalic hinge (Th) in insets, right. A region of the ganglionic eminence (GE) where Fgf8 is not expressed is shown and was used to normalise specific Fgf8 expression within the Th with background immunoreactivity as quantified in D.

(E) Phosphorylated p44/42 Mapk or Erk1/2 (p-ERK, green), and Glast-positive MZG (red) in horizontal sections of E15.5 wildtype, *Dcc*$^{kanga}$ mice and *Dcc* knockout mice reveal extent of IHF (yellow brackets) and remodelled regions of the septum (white brackets) with p-ERK-positive MZG in insets.

(F) Nuclear factor I (NFI) A or B (green), and pan-Laminin (LAM)-positive leptomeninges and basement membrane (magenta) in horizontal sections of E14 and E15 wildtype and *Dcc*$^{kanga}$ mice. NFI-positive/Glast-positive MZG cell bodies at the IHF surface are outlined with white boxes and quantified in G. Data is represented as mean ± SEM. Significant differences were determined with non-parametric Mann-Whitney tests. ns = not significant, Hi = telencephalic hinge. LAM = laminin

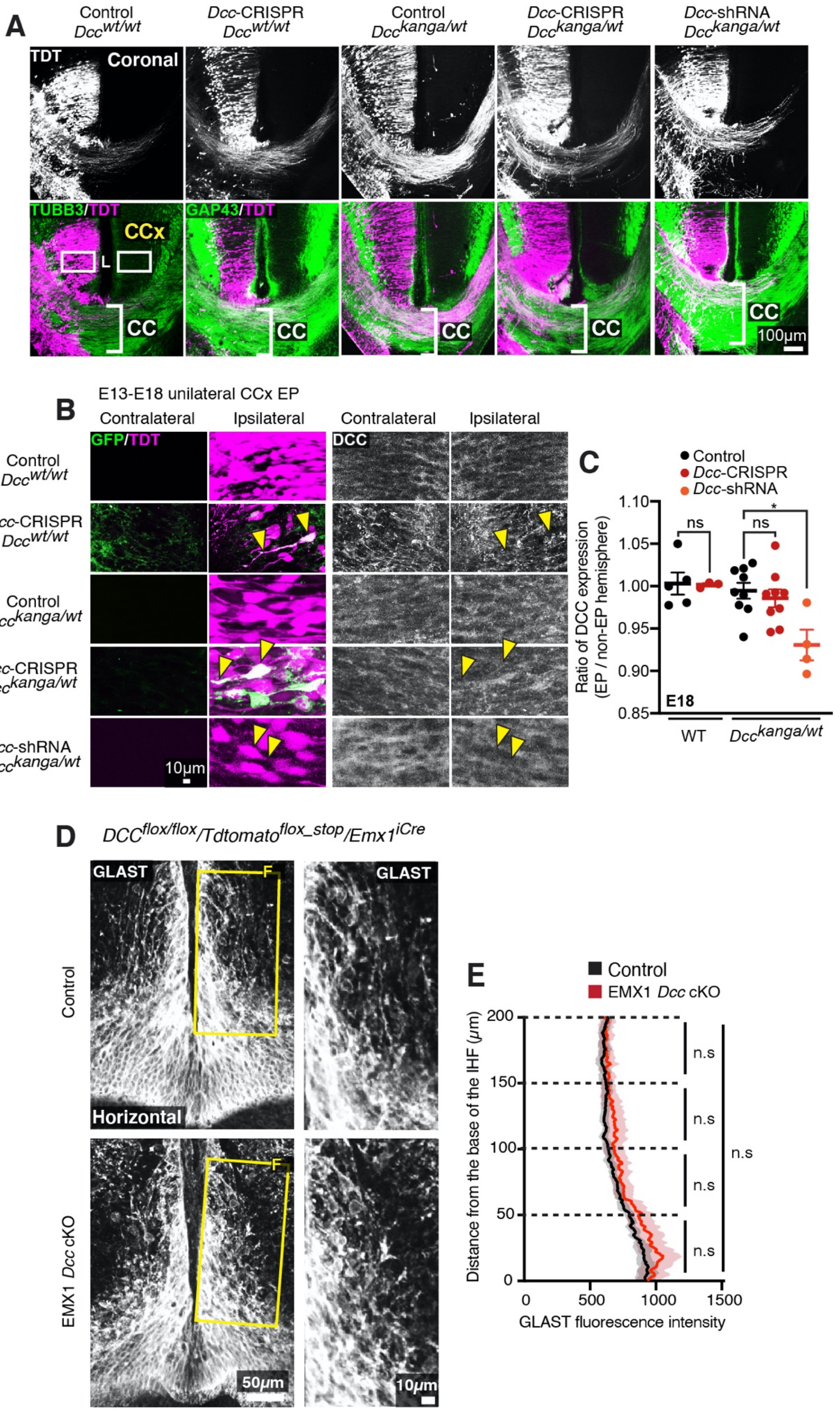
A
Control Dcc wt/wt
Dcc-CRISPR Dcc wt/wt
Control Dcc kanga/wt
Dcc-CRISPR Dcc kanga/wt
Dcc-shRNA Dcc kanga/wt
TDT
Coronal
TUBB3/TDT
GAP43/TDT
CCx
L
CC
100µm
B
E13-E18 unilateral CCx EP
Contralateral
Ipsilateral
GFP/TDT
DCC
10µm
C
Control
Dcc-CRISPR
Dcc-shRNA
ns
*
Ratio of DCC expression (EP / non-EP hemisphere)
1.10
1.05
1.00
0.95
0.90
0.85
E18
WT
Dcc kanga/wt
D
DCC flox/flox/Tdtomato flox_stop/Emx1 iCre
GLAST
F
Horizontal
Control
EMX1 Dcc cKO
50µm
10µm
E
Control
EMX1 Dcc cKO
Distance from the base of the IHF (µm)
200
150
100
50
0
n.s
0
500
1000
1500
GLAST fluorescence intensity

**Figure S6: *Dcc* knockdown via targeted in utero electroporation does not cause CC abnormalities.**

(A) TUBB3 (green) and TDT (white or magenta) in E18 *Dcc*$^{kanga}$ mice electroporated with pCAG-TDTOMATO and either *Dcc*-CRISPR or *Dcc*-shRNA constructs into the CCx at E13. The CC is outlined with white brackets and white boxes indicate the location of panels represented in B.

(B) GFP (green), TDT (magenta) or DCC (white) in E18 *Dcc*$^{kanga}$ mice electroporated with pCAG-TDTOMATO and either *Dcc*-CRISPR or *Dcc*-shRNA constructs into the CCx at E13. GFP indicates expression of the *Dcc*-CRISPR and yellow arrowheads indicate the location of select electroporated cells.

(C) Quantification of the ratio of DCC expression between ipsilateral (electroporated; EP) and contralateral (non-electroporated) hemispheres shown in B.

(D) Glial marker GLAST (white), in E15 *Dcc* cKO mice demonstrates the distribution of GLAST-positive MZG. Yellow boxes indicate region shown in insets, right and quantified in E.

(E) Quantification of mean GLAST fluorescence within the telencephalic hinge from insets in D.

All graphs represent mean ± SEM. Statistics by Mann-Whitney test or unpaired t test: *p < 0.05, n.s = not significant with p > 0.05. See related Figure 6 and Table S1.

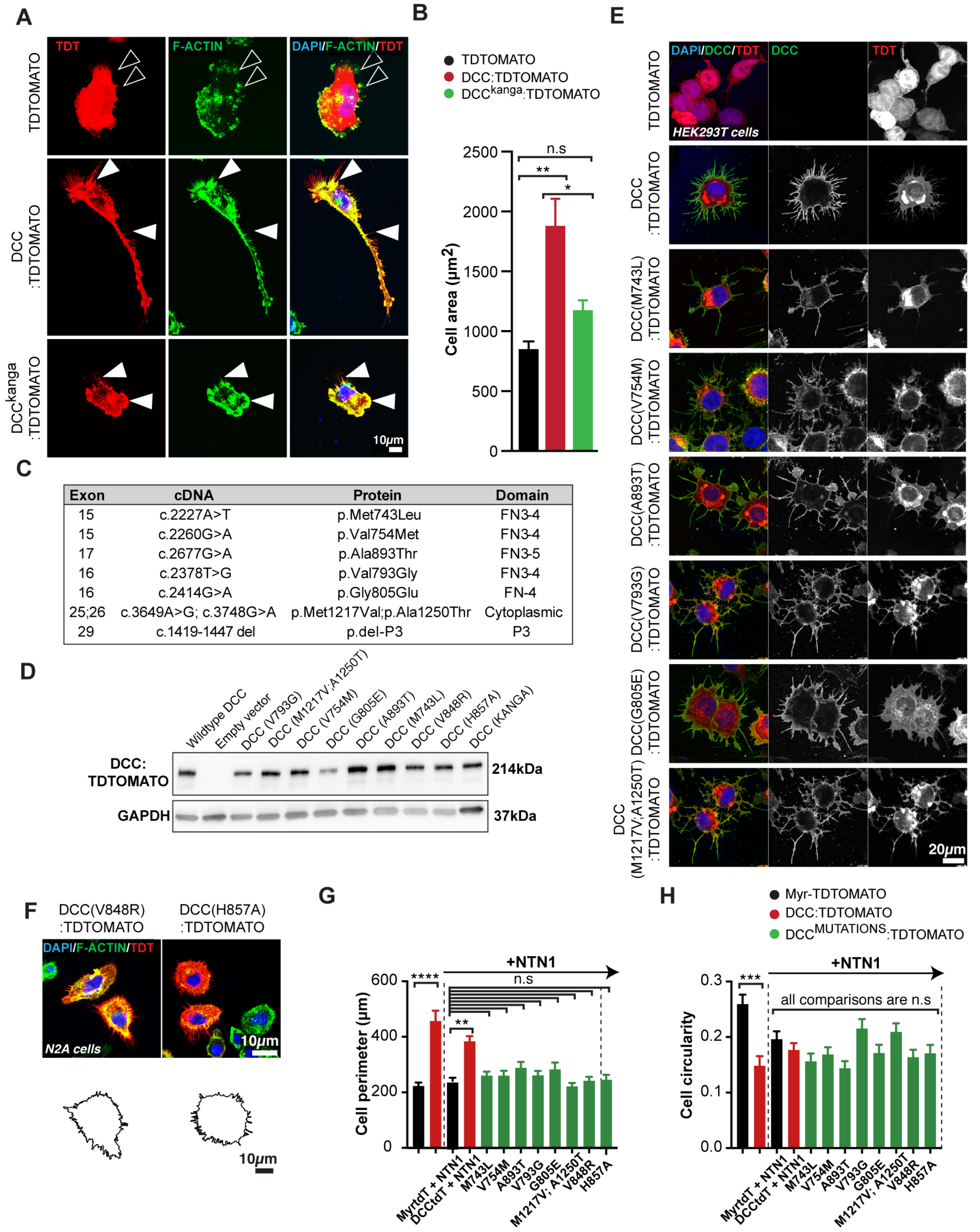


| Exon | cDNA | Protein | Domain |
|---|---|---|---|
| 15 | c.2227A>T | p.Met743Leu | FN3-4 |
| 15 | c.2260G>A | p.Val754Met | FN3-4 |
| 17 | c.2677G>A | p.Ala893Thr | FN3-5 |
| 16 | c.2378T>G | p.Val793Gly | FN3-4 |
| 16 | c.2414G>A | p.Gly805Glu | FN-4 |
| 25;26 | c.3649A>G; c.3748G>A | p.Met1217Val;p.Ala1250Thr | Cytoplasmic |
| 29 | c.1419-1447 del | p.del-P3 | P3 |

**Figure S7: Mutant *DCC* receptors are expressed and trafficked normally but are unable to modulate cell shape**

(A) Representative images of U251 glioblastoma cells immunolabelled for TDTOMATO (red), and F-actin (green) following transfection with plasmids encoding TDTOMATO, DCC:TDTOMATO, or $DCC^{kanga}$:TDTOMATO demonstrating predominant presence or absence of colocalised TDTOMATO with F-actin (arrowheads).

(B) Quantification of average cell area from U251 cells represented in A.

(C) Specific missense mutations were introduced into mouse pCAG-DCC:TDTOMATO and exon 29 was removed (del = deleted) to create the $DCC^{kanga}$:TDTOMATO construct.

(D) COS-7 cells were transfected with pCAG-DCC:TDTOMATO constructs, including those carrying specific point mutations and the $DCC^{kanga}$:TDTOMATO construct. After 48 hours, cells were lysed, and a western blot was performed for mouse DCC and GADPH. Specific bands at 214kD and 37kD are shown.

(E) HEK293T cells were transfected with pCAG-DCC:TDTOMATO constructs, including those carrying specific point mutations. After 24 hours, cells were fixed and immunohistochemistry was performed for the N-terminal of DCC without permeabilisation to detect membrane-inserted DCC.

(F) Representative images of N2A cells immunolabelled for TDTOMATO (red), and F-actin (green) following transfection with plasmids encoding DCC:TDTOMATO carrying missense mutations and stimulated with recombinant mouse NTN1 protein with cell perimeter outlined below. The cell perimeter and cell circularity of these cells, and those represented in Figure 7B are quantified in (G) and (H) respectively.

All graphs represent mean ± SEM. Statistics by Kruskal-Wallis test for multiple comparisons: n.s = not significant with $p > 0.05$, *$p < 0.05$, **$p < 0.01$, ***$p < 0.001$, ****$p < 0.0001$. See related Figure 6, Figure 7 and Table S1.

### Supplementary Table 1: Statistics

| Figure | Data compared | Number of samples | Statistical test | p value |
|---|---|---|---|---|
| 1B | Dcc knockout IHF length | n = 5 wt, n = 7 exp | Mann-Whitney test | 0.0012 |
| 1B | DCCK IHF length | n = 6 both conditions | Mann-Whitney test | 0.0022 |
| 1B | Ntn1-lacZ IHF length | n = 11 wt, n = 6 exp | Mann-Whitney test | 0.0002 |
| 1B | Dcc knockout vs. DCCK IHF length | n = 7 Dcc knockout, n = 6 DCCK | Mann-Whitney test | 0.9452 |
| 1B | Dcc knockout vs. Ntn1-lacZ IHF length | n = 7 Dcc knockout, n = 6 Ntn1-lacZ | Mann-Whitney test | 0.9452 |
| 1B | DCCK vs. Ntn1-lacZ IHF length | n = 6 both conditions | Mann-Whitney test | 0.5887 |
| 3G | E14 DCCK GLAST FI whole ROI | n = 5 exp, n = 6 wt | Mann-Whitney test | 0.1255 |
| 3G | E14 DCCK GLAST FI 0-50μm | n = 5 exp, n = 6 wt | Mann-Whitney test | 0.0043 |
| 3G | E14 DCCK GLAST FI 50-150μm | n = 5 exp, n = 6 wt | Mann-Whitney test | 0.7922 |
| 3G | E14 DCCK GLAST FI 150-200μm | n = 5 exp, n = 6 wt | Mann-Whitney test | 0.0303 |
| 3H | E15 DCCK GLAST FI whole ROI | n = 6 exp, n = 7 wt | Mann-Whitney test | 0.014 |
| 3H | E15 DCCK GLAST FI 0-50μm | n = 6 exp, n = 7 wt | Mann-Whitney test | 0.366 |
| 3H | E15 DCCK GLAST FI 50-150μm | n = 6 exp, n = 7 wt | Mann-Whitney test | 0.0082 |
| 3H | E15 DCCK GLAST FI 150-200μm | n = 6 exp, n = 7 wt | Mann-Whitney test | 0.0221 |
| 3I | E16 DCCK GLAST FI whole ROI | n = 6 exp, n = 8 wt | Mann-Whitney test | 0.9497 |
| 3I | E16 DCCK GLAST FI 0-50μm | n = 6 exp, n = 8 wt | Mann-Whitney test | 0.2824 |
| 3I | E16 DCCK GLAST FI 50-150μm | n = 6 exp, n = 8 wt | Mann-Whitney test | 0.4136 |
| 3I | E16 DCCK GLAST FI 150-200μm | n = 6 exp, n = 8 wt | Mann-Whitney test | 0.9497 |
| 3J | E14 DCCK NESTIN distribution | n = 6 both conditions | Mann-Whitney test | 0.0022 |
| 3J | E15 DCCK NESTIN distribution | n = 7 exp, n =8 wt | Mann-Whitney test | 0.0022 |
| 3J | E16 DCCK NESTIN distribution | n = 6 exp, n = 7 wt | Mann-Whitney test | 0.1014 |
| 3J | E14 vs. E15 wt NESTIN distribution | n = 6 E14, n = 8 E15 | Mann-Whitney test | 0.1419 |
| 3J | E14 vs. E15 DCCK NESTIN distribution | n = 6 E14, n = 7 E15 | Mann-Whitney test | 0.0047 |
| 3J | E15 vs. E16 wt NESTIN distribution | n = 8 E15, n = 7 E16 | Mann-Whitney test | 0.0059 |

| | | | | |
|---|---|---|---|---|
| 3J | E15 vs. E16 DCCK NESTIN distribution | n = 7 E15, n = 6 E16 | Mann-Whitney test | 0.0012 |
| 3J | E14 DCCK GLAST distribution | n = 6 both conditions | Mann-Whitney test | 0.0152 |
| 3J | E15 DCCK GLAST distribution | n = 7 exp, n =8 wt | Mann-Whitney test | 0.0012 |
| 3J | E16 DCCK GLAST distribution | n = 6 exp, n = 7 wt | Mann-Whitney test | 0.2949 |
| 3J | E14 vs. E15 wt GLAST distribution | n = 6 E14, n = 8 E15 | Mann-Whitney test | 0.0127 |
| 3J | E14 vs. E15 DCCK GLAST distribution | n = 6 E14, n = 7 E15 | Mann-Whitney test | 0.2343 |
| 3J | E15 vs. E16 wt GLAST distribution | n = 8 E15, n = 7 E16 | Mann-Whitney test | 0.0541 |
| 3J | E15 vs. E16 DCCK GLAST distribution | n = 7 E15, n = 6 E16 | Mann-Whitney test | 0.0023 |
| 4B | E14 SOX9 MZG DCCK 0-50$\mu$m | n = 8 exp, n = 12 wt | Two-way ANOVA with Sidak's multiple comparisons test | >0.9999 |
| 4B | E14 SOX9 MZG DCCK 50-100$\mu$m | n = 8 exp, n = 12 wt | Two-way ANOVA with Sidak's multiple comparisons test | 0.8565 |
| 4B | E14 SOX9 MZG DCCK 100-150$\mu$m | n = 8 exp, n = 12 wt | Two-way ANOVA with Sidak's multiple comparisons test | 0.9374 |
| 4B | E14 SOX9 MZG DCCK 150-200$\mu$m | n = 8 exp, n = 12 wt | Two-way ANOVA with Sidak's multiple comparisons test | >0.9999 |
| 4B | E14 SOX9 MZG DCCK 200-250$\mu$m | n = 8 exp, n = 12 wt | Two-way ANOVA with Sidak's multiple comparisons test | 0.9996 |
| 4D | E15 SOX9 MZG DCCK 0-50$\mu$m | n = 11 exp, n = 9 wt | Two-way ANOVA with Sidak's multiple comparisons test | 0.9275 |
| 4D | E15 SOX9 MZG DCCK 50-100$\mu$m | n = 11 exp, n = 9 wt | Two-way ANOVA with Sidak's multiple comparisons test | 0.9934 |
| 4D | E15 SOX9 MZG DCCK 100-150$\mu$m | n = 11 exp, n = 9 wt | Two-way ANOVA with Sidak's multiple comparisons test | 0.8976 |
| 4D | E15 SOX9 MZG DCCK 150-200$\mu$m | n = 11 exp, n = 9 wt | Two-way ANOVA with Sidak's multiple comparisons test | 0.4158 |

| | | | | |
|---|---|---|---|---|
| 4D | E15 SOX9 MZG DCCK 200-250μm | n = 11 exp, n = 9 wt | Two-way ANOVA with Sidak's multiple comparisons test | 0.0042 |
| 4D | E15 SOX9 MZG DCCK 250-300μm | n = 11 exp, n = 9 wt | Two-way ANOVA with Sidak's multiple comparisons test | 0.7722 |
| 4F | E16 SOX9 MZG DCCK 0-50μm | n = 6 exp, n = 7 wt | Two-way ANOVA with Sidak's multiple comparisons test | 0.026 |
| 4F | E16 SOX9 MZG DCCK 50-100μm | n = 6 exp, n = 7 wt | Two-way ANOVA with Sidak's multiple comparisons test | <0.0001 |
| 4F | E16 SOX9 MZG DCCK 100-150μm | n = 6 exp, n = 7 wt | Two-way ANOVA with Sidak's multiple comparisons test | <0.0001 |
| 4F | E16 SOX9 MZG DCCK 150-200μm | n = 6 exp, n = 7 wt | Two-way ANOVA with Sidak's multiple comparisons test | <0.0001 |
| 4F | E16 SOX9 MZG DCCK 200-250μm | n = 6 exp, n = 7 wt | Two-way ANOVA with Sidak's multiple comparisons test | 0.0012 |
| 4F | E16 SOX9 MZG DCCK 250-300μm | n = 6 exp, n = 7 wt | Two-way ANOVA with Sidak's multiple comparisons test | 0.73 |
| 4F | E16 SOX9 MZG DCCK 300-350μm | n = 6 exp, n = 7 wt | Two-way ANOVA with Sidak's multiple comparisons test | 0.5323 |
| 4F | E16 SOX9 MZG DCCK 350-400μm | n = 6 exp, n = 7 wt | Two-way ANOVA with Sidak's multiple comparisons test | 0.9664 |
| 4G | E14 SOX9 MZG DCCK TOTAL | n = 8 exp, n =12 wt | Two-way unpaired Student's t test | 0.6584 |
| 4G | E15 SOX9 MZG DCCK TOTAL | n = 11 exp, n = 9 wt | Two-way unpaired Student's t test | 0.0214 |
| 4G | E16 SOX9 MZG DCCK TOTAL | n = 6 exp, n = 7 wt | Mann-Whitney test | 0.0017 |
| 4G | E14 wt vs. E15 wt SOX9 MZG TOTAL | n = 12 E14, n = 9 E15 | Two-way unpaired Student's t test | 0.0023 |
| 4G | E14 DCCK vs. E15 DCCK SOX9 MZG TOTAL | n = 8 E14, n = 11 E15 | Two-way unpaired Student's t test | 0.5572 |

| | | | | |
|---|---|---|---|---|
| 4G | E15 wt vs. E16 wt SOX9 MZG TOTAL | n = 9 E15, n = 7 E16 | Mann-Whitney test | 0.2991 |
| 4G | E15 DCCK vs. E16 DCCK SOX9 MZG TOTAL | n = 11 E15, n = 6 E16 | Mann-Whitney test | 0.0002 |
| 5B | E17 DCCK total GFAP FI | n = 4 exp; n = 8 wt | Mann-Whitney test | >0.9999 |
| 5B | E17 Dcc knockout total GFAP FI | n = 4 exp; n = 8 wt | Mann-Whitney test | 0.9333 |
| 5B | E17 DCCK GFAP FI 0-450$\mu$m | n = 4 exp; n = 8 wt | Mann-Whitney test | 0.1091 |
| 5B | E17 DCCK GFAP FI 450-702.5$\mu$m | n = 4 exp; n = 8 wt | Mann-Whitney test | 0.0283 |
| 5B | E17 DCCK GFAP FI 702.5-902.5$\mu$m | n = 4 exp; n = 8 wt | Mann-Whitney test | 0.0040 |
| 5B | E17 Dcc knockout GFAP FI 0-450$\mu$m | n = 4 exp; n = 8 wt | Mann-Whitney test | 0.0485 |
| 5B | E17 Dcc knockout GFAP FI 450-702.5$\mu$m | n = 4 exp; n = 8 wt | Mann-Whitney test | 0.0485 |
| 5B | E17 Dcc knockout FI 702.5-902.5$\mu$m | n = 4 exp; n = 8 wt | Mann-Whitney test | 0.0040 |
| 5C | E17 Ntn1-lacZ GFAP FI | n = 6 exp; n = 3 control | Mann-Whitney test | 0.3810 |
| 5C | E17 Ntn1-lacZ GFAP FI 0-450$\mu$m | n = 6 exp; n = 3 control | Mann-Whitney test | 0.0238 |
| 5C | E17 Ntn1-lacZ GFAP FI 450-702.5$\mu$m | n = 6 exp; n = 3 control | Mann-Whitney test | 0.3810 |
| 5C | E17 Ntn1-lacZ GFAP FI 702.5-902.5$\mu$m | n = 6 exp; n = 3 control | Mann-Whitney test | 0.0238 |

| | | | | |
|---|---|---|---|---|
| 6E | Ratio IHF length P0 *Dcc* cKO mice ventral | n = 12 exp; n = 6 control | Mann-Whitney test | 0.0004 |
| 6E | Ratio IHF length P0 *Dcc* cKO mice middle | n = 12 exp; n = 6 control | Mann-Whitney test | <0.0001 |
| 6E | Ratio IHF length P0 *Dcc* cKO mice dorsal | n = 12 exp; n = 6 control | Mann-Whitney test | 0.0011 |
| 6F | Ratio CC length P0 *Dcc* cKO mice ventral | n = 12 exp; n = 6 control | Mann-Whitney test | 0.0080 |
| 6F | Ratio CC length P0 *Dcc* cKO mice middle | n = 12 exp; n = 6 control | Mann-Whitney test | 0.1861 |
| 6F | Ratio CC length P0 *Dcc* cKO mice dorsal | n = 12 exp; n = 6 control | Mann-Whitney test | 0.078 |
| 6G | Ratio CC depth P0 *Dcc* cKO mice ventral | n = 12 exp; n = 6 control | Mann-Whitney test | 0.0476 |
| 6G | Ratio CC depth P0 *Dcc* cKO mice middle | n = 12 exp; n = 6 control | Mann-Whitney test | 0.0006 |
| 6G | Ratio CC depth P0 *Dcc* cKO mice dorsal | n = 12 exp; n = 6 control | Mann-Whitney test | 0.0190 |
| 6H | Ratio HC length P0 *Dcc* cKO mice ventral | n = 12 exp; n = 6 control | Mann-Whitney test | 0.0047 |
| 6H | Ratio HC length P0 *Dcc* cKO mice middle | n = 12 exp; n = 6 control | Mann-Whitney test | 0.6495 |
| 6H | Ratio HC length P0 *Dcc* cKO mice dorsal | n = 12 exp; n = 6 control | Mann-Whitney test | 0.7559 |
| 6I | DCC expression P0 *Dcc* cKO mice cingulate cortex | n = 15 exp; n = 6 control | Mann-Whitney test | 0.0016 |

| | | | | |
|---|---|---|---|---|
| 6I | DCC expression P0 *Dcc* cKO mice intermediate zone | n = 15 exp; n = 6 control | Mann-Whitney test | <0.0001 |
| 6M | FI of DCC expression E15 *Dcc* cKO mice cingulate cortex | n = 6 exp, n = 5 control | Mann-Whitney test | 0.0173 |
| 6M | FI of DCC expression E15 *Dcc* cKO mice MZG | n = 6 exp, n = 5 control | Mann-Whitney test | 0.6623 |
| 6N | FI of GAP43 expression E15 *Dcc* cKO mice IHF surface | n = 6 exp, n = 5 control | Mann-Whitney test | 0.2468 |
| 7D | U251 CA myr-TDT vs. DCC:TDT vehicle | n = 72 wt DCC, n = 67 control | Kruskal-Wallis test with Dunn's multiple comparisons test | <0.0001 |
| 7D | U251 CA DCC:TDT vs. DCCK:TDT vehicle | n = 77 DCCK, n = 72 wt DCC | Kruskal-Wallis test with Dunn's multiple comparisons test | <0.0001 |
| 7D | U251 CA myr-TDT vs. DCCK:TDT vehicle | n = 77 DCCK, n = 67 control | Kruskal-Wallis test with Dunn's multiple comparisons test | >0.9999 |
| 7D | U251 CA myr-TDT vs. DCC:TDT NTN1 | n = 82 wt DCC, n = 65 control | Kruskal-Wallis test with Dunn's multiple comparisons test | 0.0015 |
| 7D | U251 CA DCC:TDT vs. DCCK:TDT NTN1 | n = 71 DCCK, n = 82 wt DCC | Kruskal-Wallis test with Dunn's multiple comparisons test | 0.0027 |
| 7D | U251 CA myr-TDT vs. DCCK:TDT NTN1 | n = 71 DCCK, n = 65 control | Kruskal-Wallis test with Dunn's multiple comparisons test | >0.9999 |
| 7D | U251 CA DCC:TDT vehicle vs. DCC:TDT NTN1 | n = 72 vehicle, n = 82 NTN1 | Kruskal-Wallis test with Dunn's multiple comparisons test | >0.9999 |
| 7E | U251 CP myr-TDT vs. DCC:TDT vehicle | n = 72 wt DCC, n = 68 control | Kruskal-Wallis test with Dunn's multiple comparisons test | <0.0001 |
| 7E | U251 CP DCC:TDT vs. DCCK:TDT vehicle | n = 77 DCCK, n = 72 wt DCC | Kruskal-Wallis test with Dunn's multiple comparisons test | <0.0001 |

| | | | | |
|---|---|---|---|---|
| 7E | U251 CP myr-TDT vs. DCCK:TDT vehicle | n = 77 DCCK, n = 68 control | Kruskal-Wallis test with Dunn's multiple comparisons test | >0.9999 |
| 7E | U251 CP myr-TDT vs. DCC:TDT NTN1 | n = 82 wt DCC, n = 67 control | Kruskal-Wallis test with Dunn's multiple comparisons test | <0.0001 |
| 7E | U251 CP DCC:TDT vs. DCCK:TDT NTN1 | n = 71 DCCK, n = 82 wt DCC | Kruskal-Wallis test with Dunn's multiple comparisons test | 0.0158 |
| 7E | U251 CP myr-TDT vs. DCCK:TDT NTN1 | n = 71 DCCK, n = 67 control | Kruskal-Wallis test with Dunn's multiple comparisons test | >0.9999 |
| 7E | U251 CP DCC:TDT vehicle vs. DCC:TDT NTN1 | n = 72 vehicle, n = 82 NTN1 | Kruskal-Wallis test with Dunn's multiple comparisons test | >0.9999 |
| 7F | U251 CC myr-TDT vs. DCC:TDT vehicle | n = 72 wt DCC, n = 68 control | Kruskal-Wallis test with Dunn's multiple comparisons test | 0.0006 |
| 7F | U251 CC DCC:TDT vs. DCCK:TDT vehicle | n = 77 DCCK, n = 72 wt DCC | Kruskal-Wallis test with Dunn's multiple comparisons test | <0.0001 |
| 7F | U251 CC myr-TDT vs. DCCK:TDT vehicle | n = 77 DCCK, n = 68 control | Kruskal-Wallis test with Dunn's multiple comparisons test | >0.9999 |
| 7F | U251 CC myr-TDT vs. DCC:TDT NTN1 | n = 82 wt DCC, n = 67 control | Kruskal-Wallis test with Dunn's multiple comparisons test | 0.0052 |
| 7F | U251 CC DCC:TDT vs. DCCK:TDT NTN1 | n = 71 DCCK, n = 82 wt DCC | Kruskal-Wallis test with Dunn's multiple comparisons test | >0.9999 |
| 7F | U251 CC myr-TDT vs. DCCK:TDT NTN1 | n = 71 DCCK, n = 67 control | Kruskal-Wallis test with Dunn's multiple comparisons test | >0.9999 |
| 7F | U251 CC DCC:TDT vehicle vs. DCC:TDT NTN1 | n = 72 vehicle, n = 82 NTN1 | Kruskal-Wallis test with Dunn's multiple comparisons test | >0.9999 |
| 7I | N2A CA myr-TDT vs. DCC:TDT vehicle | n = 55 wt DCC, n = 90 control | Kruskal-Wallis test with Dunn's multiple comparisons test | 0.0003 |

| | | | | |
|---|---|---|---|---|
| 8E | N2A CA myr-TDT vs. DCC:TDT NTN1 | n = 191 wt DCC, n = 100 control | Kruskal-Wallis test with Dunn's multiple comparisons test | 0.0005 |
| 8E | N2A CA myr-TDT vs. DCC(M743L):TDT NTN1 | n = 77 exp, n = 100 control | Kruskal-Wallis test with Dunn's multiple comparisons test | >0.9999 |
| 8E | N2A CA myr-TDT vs. DCC(V754M):TDT NTN1 | n = 95 exp, n = 100 control | Kruskal-Wallis test with Dunn's multiple comparisons test | >0.9999 |
| 8E | N2A CA myr-TDT vs. DCC(A893T):TDT NTN1 | n = 71 exp, n = 100 control | Kruskal-Wallis test with Dunn's multiple comparisons test | >0.9999 |
| 8E | N2A CA myr-TDT vs. DCC(V793G):TDT NTN1 | n = 86 exp, n = 100 control | Kruskal-Wallis test with Dunn's multiple comparisons test | >0.9999 |
| 8E | N2A CA myr-TDT vs. DCC(G805E):TDT NTN1 | n = 80exp, n = 100 control | Kruskal-Wallis test with Dunn's multiple comparisons test | 0.1063 |
| 8E | N2A CA myr-TDT vs. DCC(M1217V;A1250T):TDT NTN1 | n = 75 exp, n = 100 control | Kruskal-Wallis test with Dunn's multiple comparisons test | >0.9999 |
| 8E | N2A CA myr-TDT vs. DCC(V848R):TDT NTN1 | n = 92 exp, n = 100 control | Kruskal-Wallis test with Dunn's multiple comparisons test | 0.7138 |
| 8E | N2A CA myr-TDT vs. DCC(H857A):TDT NTN1 | n = 77 exp, n = 100 control | Kruskal-Wallis test with Dunn's multiple comparisons test | >0.9999 |
| S1 | Adult DCCK IHF length | n = 4 wt, n = 6 exp | Mann-Whitney test | 0.003 |
| S3D | FI of β-dystroglycan along IHF pial surface in DCCK E15 mice | n = 5 DCCK, n = 5 wt | Mann-Whitney test | 0.4206 |
| S3F | FI of APC along IHF pial surface in DCCK E15 mice | n = 5 DCCK, n = 5 wt | Mann-Whitney test | 0.0159 |
| S3F | FI of N-CADHERIN along IHF pial surface in DCCK E15 mice | n = 6 DCCK, n = 5 wt | Mann-Whitney test | 0.5368 |
| S3F | FI of β-catenin along IHF pial surface in DCCK E15 mice | n = 6 DCCK, n = 5 wt | Mann-Whitney test | 0.9307 |
| S3G | FI of NESTIN along IHF pial surface in DCCK E15 mice | n = 6 DCCK, n = 5 wt | Mann-Whitney test | 0.6623 |
| S4C | E12-E13 EdU+/Ki67+ MZG DCCK | n= 6 DCCK, n = 7 wt | Mann-Whitney test | 0.4219 |

| | | | | |
|---|---|---|---|---|
| S4C | E13-E14 EdU+/Ki67+ MZG DCCK | n= 8 DCCK, n = 6 wt | Mann-Whitney test | 0.1512 |
| S4C | E14-E15 EdU+/Ki67+ MZG DCCK | n= 8 DCCK, n = 6wt | Mann-Whitney test | 0.0939 |
| S4D | E12-E13 EdU+/Ki67- MZG DCCK | n= 6 DCCK, n = 7 wt | Mann-Whitney test | 0.6503 |
| S4D | E13-E14 EdU+/Ki67- MZG DCCK | n= 8 DCCK, n = 6 wt | Mann-Whitney test | 0.1419 |
| S4D | E14-E15 EdU+/Ki67- MZG DCCK | n= 8 DCCK, n = 6 wt | Mann-Whitney test | 0.8019 |
| S4F | E13 Cleaved-CASPASE3+ cells DCCK | n=4 DCCK, n = 5 wt | Mann-Whitney test | 0.9524 |
| S4F | E14 Cleaved-CASPASE3+ cells DCCK | n = 7 both conditions | Mann-Whitney test | 0.0816 |
| S4F | E15 Cleaved-CASPASE3+ cells DCCK | n = 8 both conditions | Mann-Whitney test | 0.5611 |
| S4I | E16 SOX9 IGGDCCK TOTAL | n = 5 exp, n = 6 wt | Mann-Whitney test | 0.0043 |
| S5A | E15 DCCK MZG GFAP FI | n = 6 exp, n = 6 wt | Mann-Whitney test | 0.5887 |
| S5D | E15 DCCK Fgf8 mRNA chromogenic intensity ratio | n = 6 exp, n = 3 wt | Mann-Whitney test | >0.9999 |
| S5D | E15 DCCK Mmp-2 mRNA chromogenic intensity ratio | n = 5 exp, n = 4 wt | Mann-Whitney test | 0.1111 |
| S5G | E14 DCCK NFIA MZG | n = 8 exp, n = 6 wt | Mann-Whitney test | 0.8691 |
| S5G | E14 DCCK NFIB MZG | n = 9 exp, n = 5 wt | Mann-Whitney test | 0.8227 |
| S6C | Ratio DCC expression EP/non EP hemisphere wildtype mice *Dcc-CRISPR* | n = 3 exp, n = 5 wt | Mann-Whitney test | >0.9999 |
| S6C | Ratio DCC expression EP/non EP hemisphere DCCK heterozygous mice *Dcc-CRISPR* | n = 9 exp, n =9 wt | Unpaired t test | 0.5412 |
| S6C | Ratio DCC expression EP/non EP hemisphere DCCK heterozygous mice *Dcc-shRNA* | n = 4 exp, n =9 wt | Mann-Whitney test | 0.0196 |

| | | | | |
|---|---|---|---|---|
| S6E | FI of GLAST along IHF EMX1 *Dcc* cKO total | n = 6 exp, n = 5 control | Mann-Whitney test | >0.9999 |
| S6E | FI of GLAST along IHF EMX1 *Dcc* cKO 0-50$\mu$m | n = 6 exp, n = 5 control | Mann-Whitney test | >0.9999 |
| S6E | FI of GLAST along IHF EMX1 *Dcc* cKO 50-100$\mu$m | n = 6 exp, n = 5 control | Mann-Whitney test | 0.7922 |
| S6E | FI of GLAST along IHF EMX1 *Dcc* cKO 100-150$\mu$m | n = 6 exp, n = 5 control | Mann-Whitney test | 0.7922 |
| S6E | FI of GLAST along IHF EMX1 *Dcc* cKO 150-200$\mu$m | n = 6 exp, n = 5 control | Mann-Whitney test | 0.7922 |
| S7B | U251 CA TDT vs. DCC:TDT | n = 49 wt DCC, n = 57 control | Kruskal-Wallis test with Dunn's multiple comparisons test | <0.0001 |
| S7B | U251 CA DCC:TDT vs. DCCK:TDT | n = 49 wt DCC, n = 59 DCCK | Kruskal-Wallis test with Dunn's multiple comparisons test | 0.0082 |
| S7B | U251 CA TDT vs. DCCK:TDT | n = 49 wt DCC, n = 59 DCCK | Kruskal-Wallis test with Dunn's multiple comparisons test | 0.0145 |
| S7G | N2A CP myr-TDT vs. DCC:TDT vehicle | n = 55 wt DCC, n = 90 control | Kruskal-Wallis test with Dunn's multiple comparisons test | <0.0001 |
| S7G | N2A CP myr-TDT vs. DCC:TDT NTN1 | n= 191 wt DCC, n = 100 control | Kruskal-Wallis test with Dunn's multiple comparisons test | 0.0014 |
| S7G | N2A CP myr-TDT vs. DCC(M743L):TDT NTN1 | n = 77 exp, n = 100 control | Kruskal-Wallis test with Dunn's multiple comparisons test | >0.9999 |
| S7G | N2A CP myr-TDT vs. DCC(V754M):TDT NTN1 | n = 95 exp, n = 100 control | Kruskal-Wallis test with Dunn's multiple comparisons test | >0.9999 |
| S7G | N2A CP myr-TDT vs. DCC(A893T):TDT NTN1 | n = 71 exp, n = 100 control | Kruskal-Wallis test with Dunn's multiple comparisons test | >0.9999 |

| | | | | |
|---|---|---|---|---|
| S7G | N2A CP myr-TDT vs. DCC(V793G):TDT NTN1 | n = 86 exp, n = 100 control | Kruskal-Wallis test with Dunn's multiple comparisons test | >0.9999 |
| S7G | N2A CP myr-TDT vs. DCC(G805E):TDT NTN1 | n = 80exp, n = 100 control | Kruskal-Wallis test with Dunn's multiple comparisons test | >0.9999 |
| S7G | N2A CP myr-TDT vs. DCC(M1217V;A1250T):TDT NTN1 | n = 75 exp, n = 100 control | Kruskal-Wallis test with Dunn's multiple comparisons test | >0.9999 |
| S7G | N2A CP myr-TDT vs. DCC(V848R):TDT NTN1 | n = 92 exp, n = 100 control | Kruskal-Wallis test with Dunn's multiple comparisons test | >0.9999 |
| S7G | N2A CP myr-TDT vs. DCC(H857A):TDT NTN1 | n = 77 exp, n = 100 control | Kruskal-Wallis test with Dunn's multiple comparisons test | >0.9999 |
| S7G | N2A CP DCC:TDT vehicle vs. NTN1 | n= 191 wt DCC vehicle, n = 100 NTN1 | Kruskal-Wallis test with Dunn's multiple comparisons test | >0.9999 |
| S7H | N2A CC myr-TDT vs. DCC:TDT vehicle | n = 55 wt DCC, n = 90 control | Kruskal-Wallis test with Dunn's multiple comparisons test | 0.0001 |
| S7H | N2A CC myr-TDT vs. DCC:TDT NTN1 | n= 191 wt DCC, n = 100 control | Kruskal-Wallis test with Dunn's multiple comparisons test | 0.3311 |
| S7H | N2A CC myr-TDT vs. DCC(M743L):TDT NTN1 | n = 77 exp, n = 100 control | Kruskal-Wallis test with Dunn's multiple comparisons test | >0.9999 |
| S7H | N2A CC myr-TDT vs. DCC(V754M):TDT NTN1 | n = 95 exp, n = 100 control | Kruskal-Wallis test with Dunn's multiple comparisons test | >0.9999 |
| S7H | N2A CC myr-TDT vs. DCC(A893T):TDT NTN1 | n = 71 exp, n = 100 control | Kruskal-Wallis test with Dunn's multiple comparisons test | 0.5619 |
| S7H | N2A CC myr-TDT vs. DCC(V793G):TDT NTN1 | n = 86 exp, n = 100 control | Kruskal-Wallis test with Dunn's multiple comparisons test | >0.9999 |
| S7H | N2A CC myr-TDT vs. DCC(G805E):TDT NTN1 | n = 80exp, n = 100 control | Kruskal-Wallis test with Dunn's multiple comparisons test | >0.9999 |

| | | | | |
|---|---|---|---|---|
| S7H | N2A CC myr-TDT vs. DCC(M1217V;A1250T):TDT NTN1 | n = 75 exp, n = 100 control | Kruskal-Wallis test with Dunn's multiple comparisons test | >0.9999 |
| S7H | N2A CC myr-TDT vs. DCC(V848R):TDT NTN1 | n = 92 exp, n = 100 control | Kruskal-Wallis test with Dunn's multiple comparisons test | >0.9999 |
| S7H | N2A CC myr-TDT vs. DCC(H857A):TDT NTN1 | n = 77 exp, n = 100 control | Kruskal-Wallis test with Dunn's multiple comparisons test | >0.9999 |
| S7H | N2A CC DCC:TDT vehicle vs. NTN1 | n= 191 wt DCC vehicle, n = 100 NTN1 | Kruskal-Wallis test with Dunn's multiple comparisons test | >0.9999 |

CA = cell area, CP = cell perimeter, DCCK = DCCKanga, E = embryonic day, EP = electroporated, exp = experimental, FI = fluorescence intensity, IGG = indusium griseum glia, MZG = midline zipper glia, P = postnatal day, ROI = region of interest, TDT = TDTOMATO, vs. = versus, wt = wildtype.
